# Molecular and Cellular Determinants of Human Iron Overload Cardiomyopathy

**DOI:** 10.64898/2026.02.02.703307

**Authors:** Sayli S. Modak, Lina Greenberg, W. Tom Stump, Akiva E. Greenberg, Nathaniel Huebsch, Michael J. Greenberg

**Affiliations:** Department of Biochemistry and Molecular Biophysics, Washington University School of Medicine, St. Louis, MO, 63110, USA; Department of Biomedical Engineering, McKelvey School of Engineering, Washington University in St. Louis, St. Louis, MO, 63130, USA

**Keywords:** Iron overload cardiomyopathy, contractility, cardiomyocytes, cardiac fibroblasts

## Abstract

Iron overload cardiomyopathy (IOC) is a serious heart condition that is caused by elevated levels of systemic iron. IOC is characterized by both systolic and diastolic dysfunction as well as arrhythmias. It has been challenging to isolate the cardiac-specific cellular and molecular mechanisms driving IOC because the disease affects multiple interconnected organ systems. Here, we leverage stem cell technologies, cardiac tissue engineering, and protein reconstitution assays to model key aspects of human IOC in vitro and to probe the cellular and molecular mechanisms driving cardiac dysfunction. We demonstrate that human engineered heart tissues consisting of both cardiomyocytes and cardiac fibroblasts faithfully recapitulate key aspects of the human disease, including reduced systolic function, impaired diastolic function, and increased prevalence of arrhythmogenic events. We demonstrate that while both cardiomyocytes and cardiac fibroblasts show increased intracellular iron levels, leading to reduced viability, cardiomyocytes show higher levels of iron accumulation and higher levels of reactive oxygen species production. Moreover, we show that in a tissue, iron overload has little effect on the action potential kinetics; however, it directly impacts the amplitude and kinetics of the calcium transient, potentially driving arrhythmogenesis. Finally, we demonstrate that iron overload decreases force production, in part, through oxidative damage of sarcomeric proteins and direct iron-based inhibition of myosin. In summary, our results reveal new insights into the cellular and molecular mechanisms of human IOC pathogenesis, and they establish new in vitro models that can be harnessed to faithfully recapitulate key aspects of the human disease phenotype.

**Highlights:**

- Contractile aspects of iron overload cardiomyopathy have been difficult to study in vitro.
- We developed engineered heart tissues to model key aspects of the human disease.
- In vitro iron overload reduces contractility and induces arrhythmogenesis.
- Iron differentially affects cardiomyocytes and cardiac fibroblasts.
- Iron overload directly impacts the actomyosin contractile apparatus.

## Introduction

Iron is essential for multiple cellular and physiological processes, ranging from oxygen transport to acting as a co-factor in enzymatic reactions; however, its systemic levels need to be tightly regulated since too little iron causes anemia while too much iron can cause damage to multiple organs, including the heart. As such, there are multiple feedback loops to regulate the amount of available iron both locally and systemically [1]. Disruption of these complex regulatory pathways can cause diseases, including cardiomyopathy [2–5].

Iron overload cardiomyopathy (IOC) is a serious form of heart disease characterized by both systolic and diastolic dysfunction, and it is often accompanied by arrhythmias [6–8]. If left untreated, IOC can cause heart failure and become a life-threatening condition. While IOC can be caused by mutations in the proteins involved in iron handling, it is most frequently seen in patients that require multiple blood transfusions, such as those with sickle cell anemia or beta-thalassemia [5–9].

It has been challenging to model and mechanistically study key aspects of the cardiac manifestations of human IOC due to several factors [10–12]. Much of our understanding of iron homeostasis comes from excellent studies in model organisms; however, there are important physiological differences in iron handling between species that can result in phenotypes that differ from the human phenotype [13–15]. Moreover, it has been difficult to isolate cardiac-specific regulation of iron from systemic regulatory mechanisms [10]. Finally, it has been difficult to mechanistically dissect the roles of specific cellular and molecular factors driving the disease progression in a humanized system [16,17]. Taken together, new tools are needed that can faithfully recapitulate key aspects of human IOC for mechanistic and translational studies.

Recent advances in stem cell technologies and tissue engineering have enabled the study of human cardiac tissues in vitro [18–28]. Importantly, these systems can be completely defined in composition, enabling the study of human cardiac-specific mechanisms that can be obscured by the complexity of systemic regulatory pathways [29,30]. Moreover, there are many cell types in the human heart, and these engineered human heart tissues can capture key aspects of crosstalk between cell types in the heart [31]. Here, we leveraged these technologies to test whether it is possible to model key aspects of human IOC in vitro and to dissect the roles of cellular and molecular factors in disease pathogenesis. Our results provide new insights into the cardiac-specific mechanisms underlying human IOC.

## Results

Our goal was to better understand the molecular and cellular bases of human IOC. As a critical first step, we needed to test whether it was possible to faithfully recapitulate key aspects of human IOC on cardiac contractility in a completely defined engineered heart tissue (EHT). Specifically, we wanted to test whether feeding iron to EHTs through media is sufficient to reproduce the reduced systolic function, impaired diastolic function, and arrhythmogenesis seen in patients. There are many cell types in the healthy and failing human heart [31], and it is not possible to include all of them in our minimalist system. Therefore, we focused on cardiomyocytes and cardiac fibroblasts which have been shown to play key roles in systolic and diastolic function as well as action potential propagation [32–34].

### Optimizing conditions to mimic iron overload

Human stem cell derived cardiomyocytes (CMs) and cardiac fibroblasts (CFs) were derived from the same parent line using well-established differentiation protocols based on small molecule modulation of the WNT-pathway as we have previously done (**Fig. S1**) [35]. Chronic iron overload can eventually lead to cell death, but this typically appears late in the disease stage due to long-term accumulation of iron [6,7]. As such, we set out to find an appropriate dose of iron for treating cells that could induce cellular dysfunction while maintaining viability.

We exposed CMs and CFs to a range of iron concentrations from 0-500 µM for 5 days (**Fig. 1**). Cells were stained to identify viable (Hoechst 33342 and Calcein-AM) and dead cells (EthD-1) (**Figs. 1a** and **S2**). Two independent markers of viability were used to ensure robust results in the event of iron-induced quenching of fluorescence. We observed a dose-dependent increase in cell death with increasing iron concentrations in both CMs and CFs (2-way ANOVA, CMs: p < 0.0001, CFs: p < 0.0001). We saw that treatment of cells with 100 µM iron had no significant effect on viability (**Figs. 1b** and **1c**) for either CMs (0 µM iron: 94 (−9/+5)% viable vs 100 µM iron: 91 (−8/+4)% viable, p = 0.66) or CFs (0 µM iron 96 (−5/+4)% viable vs 100 µM iron: 93 (−14/+7)% viable, p = 0.64); however, the percent viability significantly decreased at higher iron concentrations compared to control. This result is consistent with a previous study looking at the viability of stem cell derived cardiomyocytes at a similar iron concentration [30].

**Figure 1:**
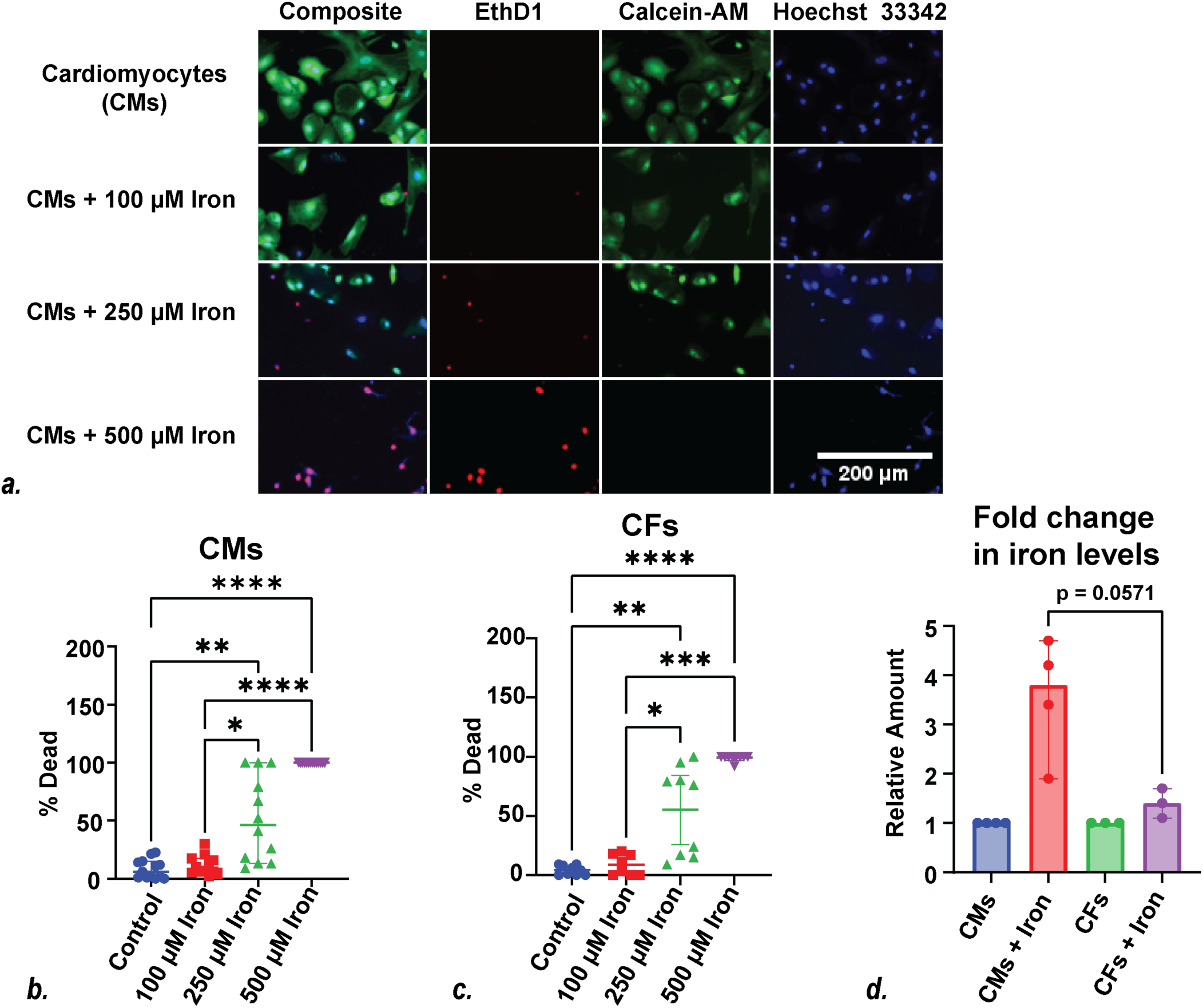
Viability and amount of iron accumulation within cardiomyocytes (CMs) and cardiac fibroblasts (CFs) in monolayer culture. (a.) Representative images of CMs survival analyzed by cell staining with Calcein-AM (green), Hoechst 33343 (blue) and EthD1 (red) with different concentrations of iron (0, 100, 250, 500 μM) fed through media for 5 days. (b. and c.) Quantification of the percent dead CMs and CFs after 5 days of treatment with iron. No significant differences were seen between control (CMs: n=13 fields of view, CFs: n=9) and 100 μM iron treated cells (CMs: n=12, CFs: n=8) while cells treated with 250 μM iron (CMs: n=12, CFs: n=9) or 500 μM iron (CMs: n=14, CFs: n=9) showed decreased viability. Data analyzed by one-way ANOVA, uncorrected Dunn’s multiple comparisons test. (d.) Colorimetric measurement of the relative levels of intracellular iron accumulation within CMs and CFs after exposure to 100 μM iron for 5 days. The amount of iron in CMs is three-fold higher than CFs when treated with the same concentration of iron (Mann-Whitney tests). Each data set represents at least two distinct differentiations or biological replicates, and there are at least three sample replicates per differentiation. Data are median ± 95% CI, *p-value≤0.05, **p-value≤0.01, ***p-value≤0.001, and ****p-value≤0.0001.

We also wanted to ensure that feeding 100 µM iron for 5 days would cause an appreciable increase in intracellular iron. We used a colorimetric assay to quantify the amount of labile iron and observed that treatment of CMs with 100 µM iron for 5 days (**Fig. 1d**) caused a ∼3.5-fold increase of intracellular iron (p=0.03). This is within the range of fold changes observed in patients; however, it should be noted that there is large variability between patients, since chronic iron exposure causes accumulation of iron over time [36–40]. In contrast, iron treatment did not cause a statistically significant change in CFs (p = 0.7). Given these results, we treated tissues with 100 µM iron for 5 days in all further experiments.

### Iron treatment of engineered heart tissues consisting of CFs and CMs is sufficient to recapitulate key features of human IOC

Key features of IOC in patients are decreased systolic function, impaired diastolic function, and arrhythmias. We wanted to test whether iron treatment of EHTs composed of CFs and CMs is sufficient to capture these key contractile aspects of the human phenotype. To investigate this possibility, we used two complementary EHT systems: one tailored for mechanical measurements of contractility and one tailored to fluorescence-based measurements of action potentials and calcium transients.

To examine how iron affects tissue contractility, we generated ring-shaped tissues composed of both CMs and CFs in a collagen-Matrigel matrix [19,41]. Tissues were grown under passive tension for 14 days and treated with 100 µM iron for 5 days before mechanical testing. The ring-shaped tissues were then mounted between a length mover and a force transducer for mechanical testing. Each tissue was subjected to a series of stretches under 1 Hz electrical stimulation and the total force exerted by the tissue was measured (**Fig. S3**), where the total force is the sum of the active force generated by beating cardiomyocytes and the passive force resisting stretch exerted by cells in the tissue (e.g., fibroblasts) and the matrix.

The active force, which is related to systolic function (i.e., the force produced during heart contraction), was measured as the force above baseline produced by cardiomyocyte beating (**Fig. 2a**). We observed that iron treatment significantly reduces the active forces produced by the tissue (**Fig. 2b**; p < 0.0001), consistent with the decreased systolic function seen in patients. Moreover, we observed that both iron treated and control tissues showed increases in active force as the tissue was stretched to longer lengths, an effect related to the Frank-Starling response; however, this response was blunted in iron-treated tissues (**Fig. 2b**). We also saw that while iron treatment did not change the amount of time required for contraction or relaxation (**Figs. S4** and **S5**), it did cause significant reductions in the speeds of both contraction and relaxation (p < 0.0001 and p < 0.0001, respectively) (**Figs. S4** and **S6**). The passive force, which is related to diastolic function (i.e., passive stretching during filling of the heart), was the baseline force measured at each length (**Figs. 2a** and **S3**). The passive forces observed at longer stretches were higher for the iron-treated tissues than the control tissues (**Fig. 2c**). This is consistent with diastolic dysfunction seen in patients.

**Figure 2:**
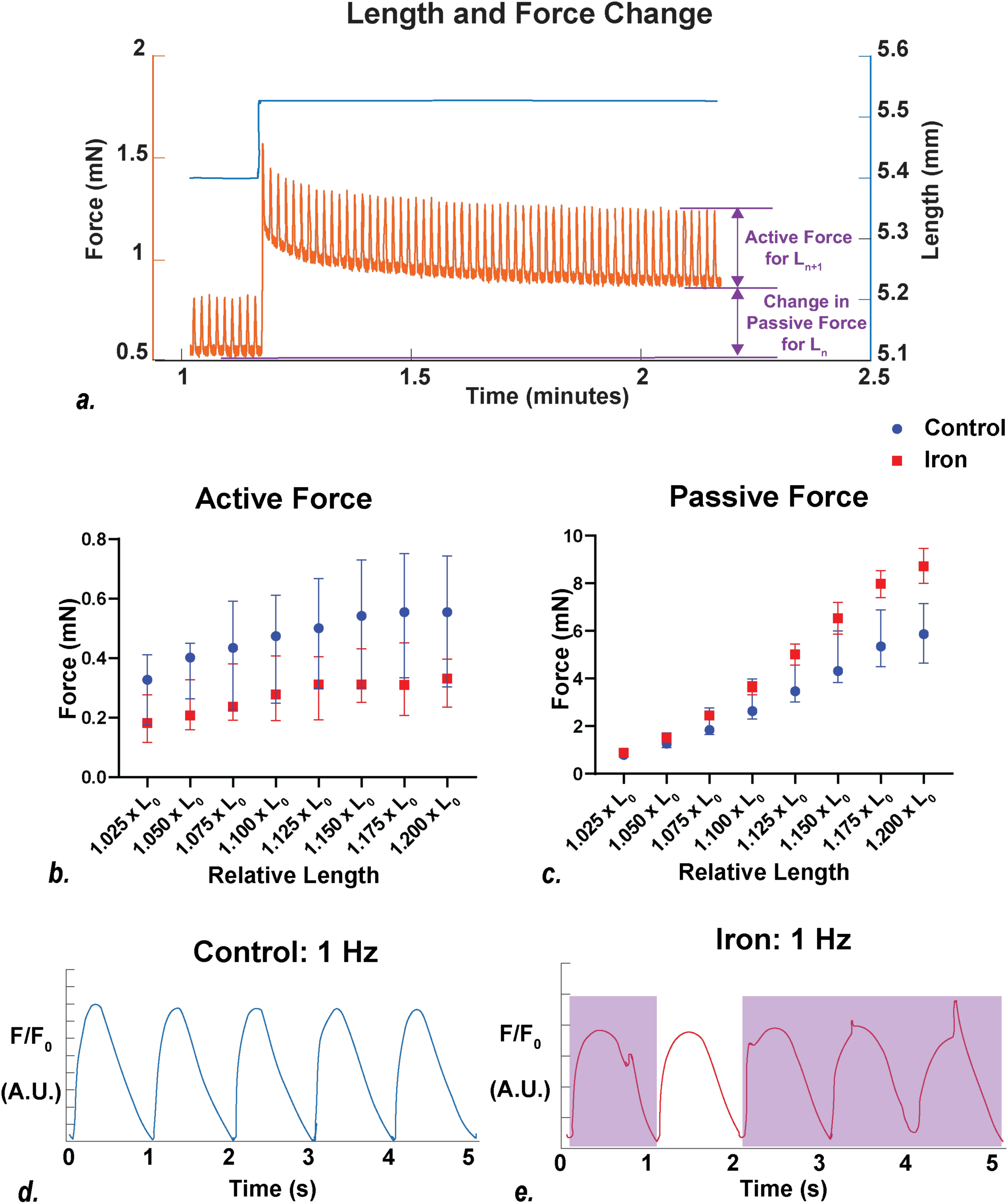
EHTs capture key contractile aspects of iron overload cardiomyopathy. (a.) Representative data trace showing the force produced by the EHT (orange) as a function of time in response to a length step (blue). Traces were used to calculate the active and passive forces. (b.) Active force and (c.) passive force exerted by the tissue as a function of length. n = 24 control tissues (blue) and n = 25 iron-treated tissues (red). Statistical testing was done by two-way ANOVA. Each data set represents three or more distinct differentiations or biological replicates. Data are median ± 95% CI, *p-value≤0.05, **p-value≤0.01, ***p-value≤0.01, and ****p-value≤0.01. Representative traces showing the fluorescence of the calcium sensitive dye as a function of time. EHTs were stimulated at 1 Hz. Traces for (d.) control and (e.) iron-treated EHTs are shown. Irregular waveforms are highlighted by purple boxes. Additional traces and quantification can be found in the supplementary materials.

Finally, we generated ‘dog-bone’ shaped EHTs known as micro–Heart Muscles (μHMs) [28] (**Fig. 3a**) that were optimized for high-resolution optical imaging to test whether they display characteristics of arrhythmogenesis such as altered calcium handling [42]. μHMs were grown under uniaxial tension for 14 days and treated with iron for 5 days. The tissues were then loaded with the calcium-sensitive FluoForte dye before optically imaging under electrical stimulation. Both iron treated and control EHTs responded to electrical stimulation, causing transient increases and decreases in fluorescence (**Figs. 2d** and **2e**). Importantly, iron-treated tissues showed a higher frequency of aberrant calcium transients (**Fig. 2e**), consistent with a higher propensity for arrhythmogenesis. Compared to control tissues that generally exhibited regular waveforms, iron-treated EHTs frequently showed distorted waveforms with spikes in intensity. Examples of transients collected at 1, 1.5, and 2 Hz can be found in (**Figs. S7, S8,** and **S9**). We observed that iron-treated tissues showed higher propensity for abnormal calcium transients at 1 and 1.5 Hz stimulation compared to the control, and that both control and iron-treated tissues showed a similar percentage of abnormal transients at high stimulation frequency of 2 Hz (**Fig. S10**). Altogether, our results clearly demonstrate that iron treatment of EHTs composed of CFs and CMs is sufficient to capture key aspects of the systolic dysfunction, diastolic dysfunction, and arrhythmogenesis seen in patients.

**Figure 3:**
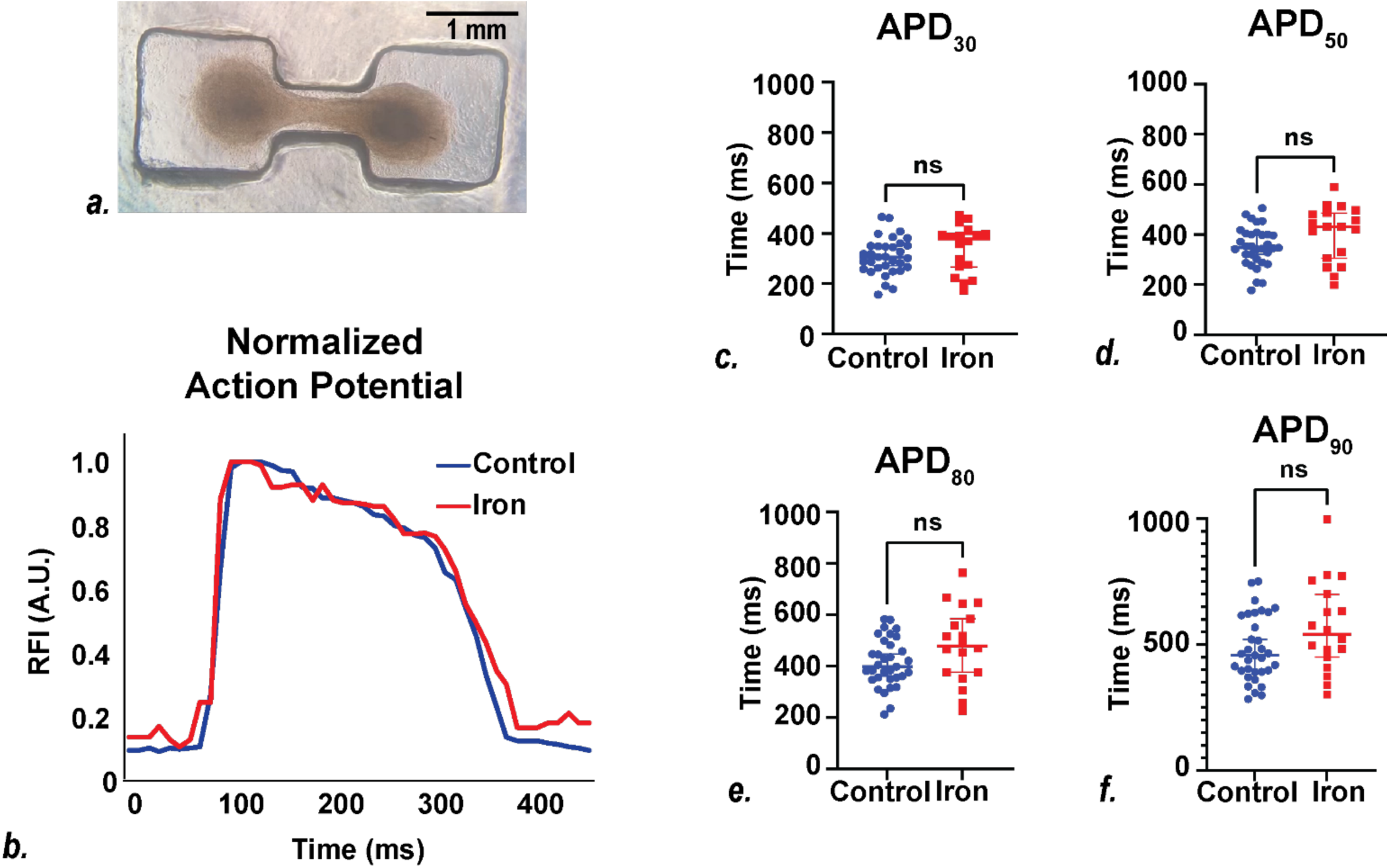
Effect of excess iron on the action potentials of engineered heart tissues. (a.) Image of a representative micro–heart muscle (μHM) engineered heart tissue formed in a dog bone shaped PDMS stencil. (b.) Representative normalized voltage waveform for a μHM. µHMs were treated with 100 µM iron for 5 days and loaded with a voltage sensitive dye to measuring action potentials by high optical resolution microscopy. Waveforms for control (blue) and iron-treated (red) μHMs. (c-f) Analysis of fluorescence waveforms to measure voltage. Shown are the action potential durations (APD) for the signal to return to 30, 50, 80, and 90% of the peak signal. Statistical testing was done using a Mann-Whitney test. Each point shows an individual tissue, the central bar shows the median, and the error bars show 95% confidence intervals. Data was drawn from three or more distinct differentiations. n = 33 for control tissues (blue) and n = 18 for iron-treated tissues (red). *p-value≤0.05, **p-value≤0.01, ***p-value≤0.01, and ****p-value≤0.01.

### Iron overload affects calcium transients without significantly affecting action potential kinetics

Given that our result that CMs and CFs are sufficient to capture key aspects of the contractile phenotype, we set out to better understand the molecular and cellular bases for these changes. Cardiac contraction is initiated by an action potential, which then causes the release of calcium, and this is followed by myosin-based sarcomeric contraction. Given that we saw impaired contraction, this could come from changes in excitation, calcium release, and/or myosin-based contraction. We therefore set out to test each of these possibilities.

The kinetics of the action potential were measured in μHMs [28] using the voltage sensitive dye, FluoVolt (**Figs. 3b** and **S11**). Both control and iron-treated tissues show clear action potential waveforms. We did not observe any statistically significant changes in the kinetics of the action potential between control and iron-treated EHTs (APD_30_: p = 0.22, APD_50_: p = 0.08, APD_80_: p = 0.09, APD_90_: p = 0.08, UPD: p=0.13, Peak time: p=0.45) (**Figs. 3c, 3d, 3e** and **3f**). Moreover, we saw a reduction in the amplitude of the action potential (**Fig. S12**). Taken together, our results suggest that in the context of an EHT, iron does not appreciably affect the kinetics of the action potential.

The action potential triggers the release of calcium in CMs. Calcium transient waveforms were measured in μHMs [28] using the calcium-sensitive FluoForte dye (**Figs. 4a, 4b and S13**). Both control and iron-treated tissues showed clear calcium transient waveforms in response to electrical stimulation. However, iron-treated tissues show a larger calcium fluorescence amplitude (p = 0.01) and larger integrated calcium flux (p = 0.01) (**Figs. 4c** and **4d**). Interestingly, the iron-treated EHTs showed prolonged calcium transients as seen by increased time to 30% relaxation (τ_30_; p = 0.0004) and time to 50% relaxation (τ_50_; p = 0.001) (**Fig. 4e, 4f** and **S14**).

**Figure 4:**
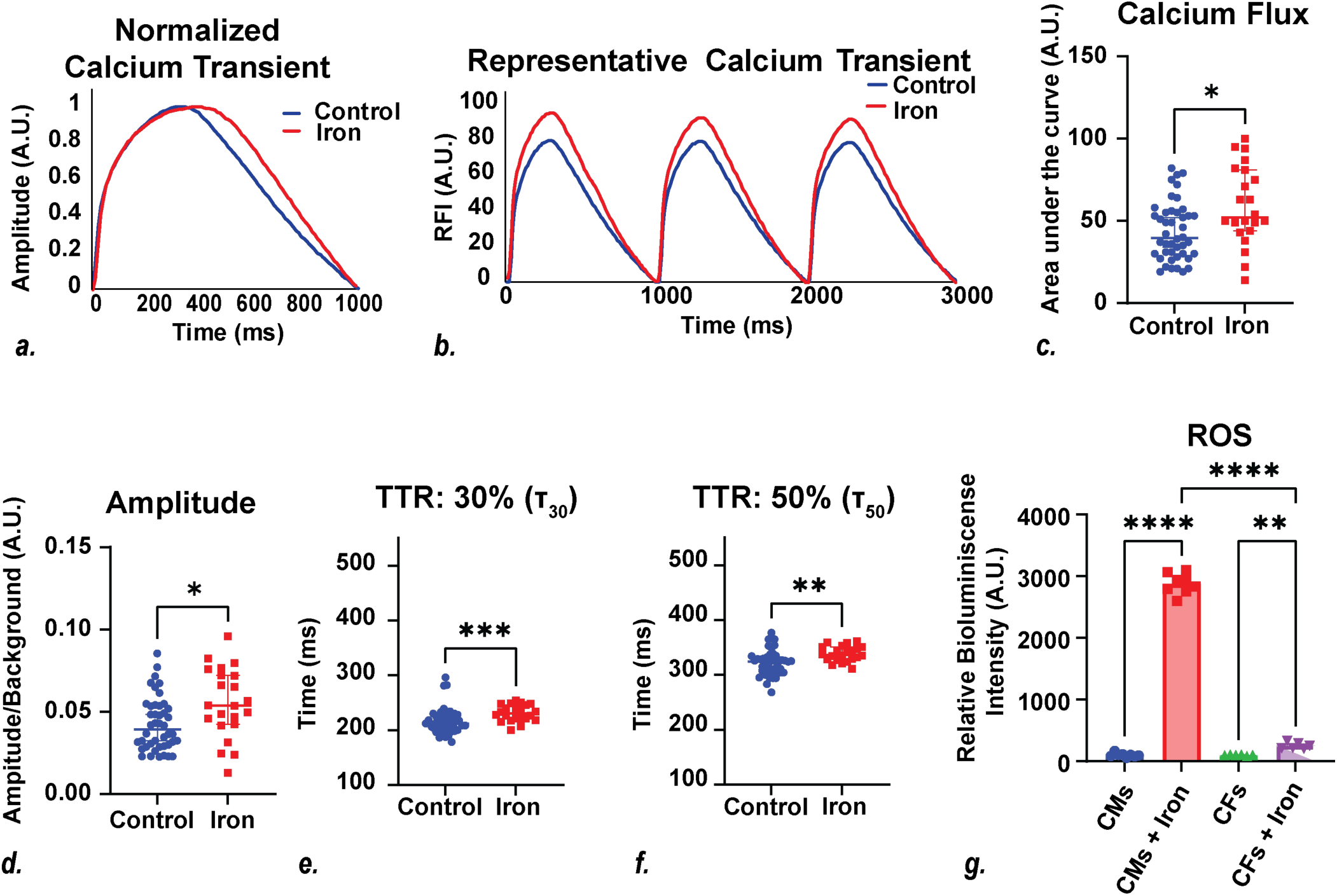
Effect of excess iron on the calcium transients of engineered heart tissues. Calcium transients were measured using calcium-sensitive dye. (a.) Representative normalized calcium transient showing the fluorescence as a function of time for a single beat of control (blue) and iron treated (red) μHMs (b.) Representative calcium transient for control (blue) and iron treated (red) μHMs. (c-f) Quantification of key parameters from calcium transients. n = 44 control (blue) and n=22 iron-fed (red) μHMs. Statistical testing was done using a Mann-Whitney test. Shown are (c.) integrated calcium flux, (d.) amplitude, (e.) Time to relax by 30% (τ_30_) and (f.) 50% (τ_50_). (g.) The amount of reactive oxygen species (ROS) was measured for both CMs and CFs using a bioluminescence assay. Wells treated with iron for 5 days (CMs: n=8, CFs: n=6) have three-fold higher ROS compared to control (CMs: n=8, CFs: n=6) as seen by Dunnett’s T multiple comparisons test. Each data point represents an individual well of a plate. Data are drawn from three or more distinct differentiations. Symbols show median and error bars show 95% confidence intervals. *p-value≤0.05, **p-value≤0.01, ***p-value≤0.01, and ****p-value≤0.01.

Changes in the calcium transient have previously been shown to be proarrhythmogenic [42]. Our observations of aberrant calcium transients (**Figs. 2d** and **2e**) and changes in the kinetics of calcium handling suggest that this may be due to iron affecting the machinery responsible for calcium handling. While a full examination of the calcium-handling pathways is beyond the scope of this study, we chose to focus on the role of reactive oxygen species (ROS). Previous studies have shown that labile iron causes the formation of ROS, and that ROS can directly affect the calcium-handling machinery [7,43–46]. Therefore, we tested the levels of ROS production in CMs and CFs using a bioluminescent luciferin assay (**Fig. 4g**). While iron treatment increased ROS production in both CMs and CFs, we found that the amount of ROS produced in CMs after iron treatment was significantly higher compared to CFs treated with same concentration of iron (p < 0.0001). This is consistent with our observation that CMs have higher levels of iron accumulation than CFs (**Fig. 1d**). Taken together, our results demonstrate changes in calcium handling associated with iron overload that could contribute to the observed arrhythmogenic phenotype seen in patients and EHTs (**Fig. 2e**).

### Iron-induced damage of the contractile apparatus contributes to systolic dysfunction

Our results show that iron treatment causes an increase in the amplitude and duration of the calcium transient (**Fig.4**) which would be expected to increase contraction; however, we observe a clear reduction in the active force production in EHTs (**Fig. 2b**). This seemingly paradoxical result suggests that iron accumulation could also affect sarcomeric-based contractility. While several mechanisms may contribute to this phenomenon, we focused on addressing whether iron treatment has direct effects on the actomyosin contractile apparatus.

To test this possibility, we purified porcine cardiac actin, which is identical to human cardiac actin, and porcine cardiac myosin, which has biophysical properties that are indistinguishable from the human isoform [47–49]. We then investigated the effects of iron treatment on actomyosin contractility using an *in vitro* motility assay in which fluorescently labeled actin filaments are propelled over a bed of myosin [50]. Importantly, this simplified assay enables the direct measurement of molecular contractility. First, we examined the effects of iron treatment on the contractile apparatus by incubating myosin and actin with 1 mM iron either overnight or for 3 days and then running the motility assay with iron included in the final buffer. We saw that treatment with 1 mM iron significantly reduced contractility after both overnight incubation and 3 days of incubation (**Fig. 5**). A similar but more modest effect was seen with incubation of 100 µM iron (i.e., the same concentration as in the cellular experiments) for 3 days (**Fig. S15**).

**Figure 5:**
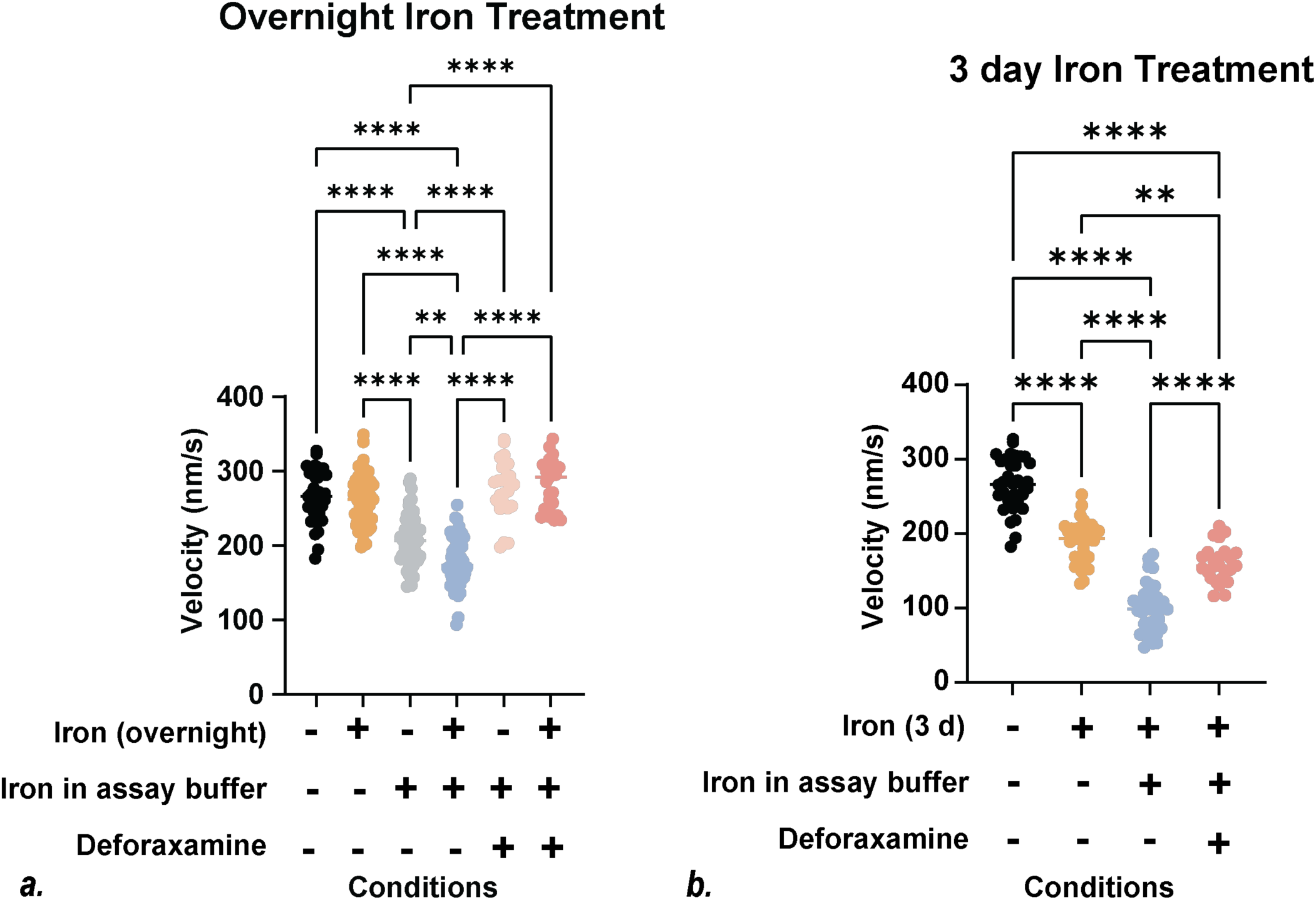
In vitro motility assays reveal iron effects on the actomyosin contractile apparatus. Motile rate of actin filaments moving over a bed of cardiac myosin. Proteins were treated with 1 mM iron (a.) overnight or (b.) 3 days. Iron was included in the buffer for some assays, and some assays contained the iron chelator deferoxamine (DFO). Each point represents an individual actin filament speed recorded over 30 seconds. Statistical testing was done using an ANOVA followed by post-hoc t-tests. *p-value≤0.05, **p-value≤0.01, ***p-value≤0.01, and ****p-value≤0.01.

The observed reduced molecular contractility could have come from either oxidative damage to actomyosin and/or from direct inhibition of myosin’s mechanochemical cycle. To distinguish between these two nonexclusive possibilities, we conducted an additional series of experiments. First, we assessed whether acute iron exposure inhibits actomyosin contractility. Here, we did not treat the actomyosin with iron before the experiment, and iron was only added in the final motility buffer. We saw that the inclusion of 1 mM iron in the final buffer directly reduces motility, demonstrating direct effects of free iron on myosin’s mechanochemical cycle (**Fig. 5a**).

Next, we evaluated whether iron causes oxidative damage of actomyosin by first incubating the actomyosin with iron for 1 or 3 days as described above but then omitting the iron from the final motility buffer. We saw that even in the absence of iron in the final buffer, pre-incubation of actomyosin with 1 mM iron overnight had no statistically significant effect on motile speed (p>0.99); however, there was a significant reduction in speed after 3 days of incubation with iron (p<0.001). This suggests that prolonged iron exposure can cause oxidative damage to the actomyosin, independent of iron in the final buffer. Interestingly, the combined effects of long-term incubation and inclusion of iron in the final buffer were greater than either factor independently (**Fig. 5**), suggesting synergistic effects of oxidative damage and direct iron-based inhibition of actomyosin motility.

Finally, we tested whether treatment with deferoxamine (DFO), an iron chelator used clinically to treat IOC patients, could improve contractility. We saw that the inclusion of DFO in the final motility buffer together with iron was sufficient to prevent iron-induced reduction in motility (**Fig. 5**). Moreover, pre-treatment with both iron and DFO for 3 days followed by inclusion of both iron and DFO in the final buffer only partially rescued the motility rate, suggesting that the DFO concentration used was unable to completely prevent oxidative damage to actomyosin. In summary, we demonstrate that the reduced contractility seen in EHTs can in part be explained by both iron-induced damage of the actomyosin contractile apparatus and iron-based direct inhibition of myosin’s biochemical cycle. Finally, we demonstrate that DFO can partially rescue motility in the presence of acute iron treatment, but that damage to the actomyosin is not reversible with chelators.

## Discussion

Here, we demonstrated that it is possible to recapitulate key aspects of human iron overload cardiomyopathy in EHTs consisting of cardiomyocytes and cardiac fibroblasts. We further demonstrated how these tools could be used to mechanistically dissect key aspects of the molecular and cellular basis of human IOC in vitro.

### Modeling of human IOC in vitro

Excellent studies using animal-based models of iron overload have revealed complex feedback mechanisms that delicately maintain iron homeostasis [10,51–54]. These complex mechanisms include both control of local iron levels in the heart as well as broader systemic regulation. Given this inherent complexity, it has been difficult to deconvolve the direct cardiac-specific effects of iron overload from systemic-based effects and secondary physiological adaptations. Moreover, there are multiple physiological differences between small animals and humans that can limit the ability of these animal models to faithfully recapitulate aspects of the human phenotype. For example, mice with a cardiac-specific ferroportin mutation that causes human IOC develop systolic dysfunction when fed a high iron diet; however, they do not develop the diastolic dysfunction that is a hallmark of IOC in patients [11].

Here, we developed an EHT system capable of recapitulating key contractile aspects of human IOC. Importantly, this system allowed us to completely define the components of the tissue, enabling us to use reductionist approaches to probe the contributions of specific cellular and molecular factors in IOC. It should be noted that our minimalist approach is not a substitute for in vivo studies since it cannot fully recapitulate the complexities of systemic iron homeostasis; rather we view our reductionist approaches as complementary to in vivo tools. However, given that this system captured key contractile aspects of IOC in vitro, we believe that this platform could potentially be harnessed for translational studies.

The EHTs incorporating stem cell derived CMs and CFs provides the cells with critical cues that promote their maturation including electrical coupling between cells, 3D interactions both between cells and the matrix, and mechanical tension [55–57]. Previous studies of other disease-causing pathologies using single cells and engineered heart tissues have demonstrated that some aspects of the disease phenotype are only unmasked in the context of a tissue [22,23,30]. For example, our 3D engineered heart tissues containing CFs and CMs did not demonstrate appreciable changes in the kinetics of the action potential; however, this result contrasts with previous studies of 2D stem cell derived CM monolayers [30].

Recent studies have demonstrated the diversity of cell types in both the healthy and failing heart [31,32]; however, it is not tractable to examine the contributions of all these cells to IOC in a single study. We focused on CMs and CFs since these cell types play central roles in the relevant disease processes, namely contraction (systolic function), regulating the extracellular matrix (diastolic function), and excitation-contraction coupling (arrhythmias) [33,34]. In the future, we envision that this same platform could be used to investigate the roles of other cell types important to IOC. For example, different macrophage polarizations have unique roles in maintaining systemic iron homeostasis [58–61], and our reductionist approach could be easily adapted to study these roles.

### Iron has different effects on human cardiomyocytes and cardiac fibroblasts

Here, we observed several differences in the responses of CMs and CFs to iron overload. We saw that while both CMs and CFs take up excess iron, the total amount accumulated was higher with CMs. This is consistent with previous studies in mice showing higher accumulation of iron in cardiomyocytes [11,29,30,40,62–64]. While there are several shared pathways for iron import between CMs and CFs, it has been shown that labile iron can additionally enter CMs through L-Type calcium channels (LTCC). The high expression of LTCC in CMs may increase the iron burden in these specific cells, and our results in human cells further support this notion [12,62,64–68]. Moreover, fibroblasts express higher levels of ferritin, enabling them to tolerate higher iron loads before the accumulation of labile iron [69,70].

Accumulation of labile iron has been shown to cause the production of reactive oxygen species via Fenton reactions, which in turn can damage and/or affect the activity of critical cellular structures involved in calcium handling including the ryanodine receptor and SERCA [43,46,71] (see below). Our results clearly demonstrate that while both CMs and CFs produce reactive oxygen species in the presence of iron, the amount of ROS produced is significantly higher for CMs. This is consistent with our observation that CMs have higher accumulation of iron compared to CFs. Moreover, while CMs had increased ROS production at lower iron concentrations, both CMs and CFs had reduced viability at high iron concentrations, likely due to iron-induced ferroptosis [69,72,73]. Taken together, our studies here clearly showed differences between human cardiomyocytes and cardiac fibroblasts in their responses to iron.

### Mechanistic insights into how iron accumulation impairs cardiac contractility

Our results clearly demonstrate that iron overload causes reduced cardiomyocyte active force production (**Fig. 2b**), consistent with the systolic dysfunction seen in patients. There are multiple processes which could affect cardiac contractility, so we investigated several of these possibilities. Our results clearly demonstrate that treatment of cells with 100 µM iron for 5 days had negligible effect on cellular viability, and therefore the observed reduction in force cannot be explained by iron-induced cell death. This is consistent with the phenotype seen in patients [1,4–9]. CMs are post-mitotic and therefore not renewable; however, the IOC phenotype is typically reversible with appropriate treatment, consistent with the notion that the observed reduced contractility stems from other molecular mechanisms [6–8].

We also considered the effects of iron on excitation-contraction coupling. Our results show that in the context of an EHT, the kinetics of the action potential are not significantly affected by iron treatment, which is consistent with some previous studies looking at single cardiomyocytes isolated from animal hearts, but not others [74,75]. It is possible that these differences could stem from species-specific differences or from the incorporation of multiple cell types in our tissues, including CFs, which can modulate the action potential [33,34]. Moreover, we observed a reduction in the amplitude of the action potential, which has been seen consistently across animal models of iron overload [76]. It is worthwhile noting that while we observed a reduction in the signal of the voltage-sensitive dye with iron, we could not exclude the possibility of iron affecting fluorescence.

We also observed that iron treatment in EHTs causes an increase in the amplitude and duration of the calcium transient (**Figs. 4c** and **4d**), as well as appearance of irregular spikes in the calcium waveforms (**Figs. 2d** and **2e**). Previous studies have shown that reactive oxygen species can damage or affect the activity of key proteins involved in calcium handling including SERCA, ryanodine receptors, the sodium-calcium exchanger, and L-type calcium channels [43–45,71]. Additionally, whole transcriptome sequencing of stem cell derived cardiomyocytes showed that iron treatment increased expression of both calsequestrin and the sodium-calcium exchanger [30]. Together, these changes can affect the rates of calcium release and reuptake [42], consistent with our observation of prolongation of the calcium transient. Moreover, our results clearly demonstrate altered calcium waveforms with irregular spikes in calcium in the presence of iron (**Figs. 2d** and **2e**). Previous studies have established that ROS can trigger calcium sparks, which are in turn proarrhythmogenic, and this is consistent with our data [44]. Taken together, our results in human EHTs provide important insights into potential mechanisms driving arrhythmogenesis in human IOC [42].

In isolation, the observed increased amplitude of the calcium transient (**Fig. 4d**) would be expected to lead to increased force of contraction; however, we observe a reduction (**Fig. 2b**). This suggests that alterations in the calcium transient are not responsible for the observed inhibition of force production. The molecular mechanisms of sarcomeric contraction involve many connected proteins [77]; however, we focused on whether iron overload can impair the force generating interactions between actin and myosin. To investigate actomyosin interactions, we reconstituted contraction in vitro using an actin filament gliding assay in which actin filaments translocate over a bed of myosin in the presence of ATP. This well-established, reductionist assay has the advantage that it retains many of the features of solution biochemistry while allowing direct measurement of actomyosin motile function [50].

Our data clearly show that iron overload impairs actomyosin interactions through at least two independent mechanisms. We saw that free iron can directly inhibit actomyosin-driven motility, and that this can be rescued by chelating free iron by deferoxamine (**Fig. 5**). As such, high iron levels can act as inhibitors of actomyosin function. Our result is consistent with a previous study showing that iron depresses contraction in isolated papillary muscles [78]. While the exact mechanism underlying this inhibition is not clear, it is well-established that metal ions play essential roles as co-factors in myosin’s ATPase cycle [79]. It is possible that iron competes with these metals or that it binds to cation binding sites on myosin; however, additional studies are needed to pinpoint this mechanism.

We also saw that exposure of actomyosin to high levels of iron over time leads to inhibition of contractility, independent of direct effects on actomyosin activity (**Fig. 2b**). This idea is supported by the observation that prolonged exposure of actomyosin to iron reduces the motile speed even in the absence of iron in the final assay buffer, and this effect is not reversible by the inclusion of chelator (**Fig. 5**). This impairment of motile function is likely due to oxidative damage of actomyosin induced by Fenton reactions. Previous studies have shown that oxidative damage can impair actomyosin interactions [80,81], and our results are consistent with oxidative damage of actomyosin playing a key role in the reduced contractility seen in IOC.

The dual mechanisms underlying iron’s effects on actomyosin have important implications for the treatment of IOC. Two treatments that are being investigated for IOC are calcium channel blockers and iron chelators. As mentioned earlier, it has been proposed that iron accumulation in CMs is promoted by calcium import through L-type calcium channels [12,62,64–68]. Based on excellent studies in mice, a clinical trial investigating whether the L-type calcium channel blocker verapamil could help patients with iron overload cardiomyopathy was initiated [67]. Moreover, there are clinical trials looking at the efficacy of iron chelation by DFO in the context of IOC. Our results demonstrate that DFO treatment, can help to alleviate acute levels of iron overload by chelating free iron that would interfere with actomyosin contraction. Interestingly, DFO treatment could not completely rescue the oxidative effects of long-term exposure to iron. This result implies that recovery from iron overload would require the turnover of damaged actomyosin in cardiomyocytes. Our results support the premise of recent clinical trials evaluating the combined effects of iron chelation and LTCC blockers [82–86].

### Limitations

Our studies are not without limitations. First, EHTs are simplified reductionist tools to study the cardiac-specific effects of IOC on contractility, and therefore, they cannot capture many critical aspects of iron overload in vivo, including systemic iron regulation and cross talk between organ systems. Moreover, our current system does not recapitulate the diversity of cells in either healthy or failing hearts. While incorporation of cells into EHTs promotes maturation, they are still never as mature as adult cardiomyocytes, and EHT systems cannot recapitulate the complex remodeling that accompanies heart failure. It is also difficult to directly compare the levels of iron seen in our cells to levels in patients, since myocardial iron levels are highly variable between patients, and the levels of myocardial iron are rarely measured outside the context of end-stage heart failure [6,8,38,39]. Moreover, iron accumulation is cumulative, and chronic iron overload over 20 years without chelation therapy can cause accumulation of very high levels of myocardial iron [39,40]. As such, our studies cannot model the full time-course of the disease. Finally, we demonstrated several potential mechanisms underlying changes in calcium handling and contractility; however, there are likely additional mechanisms that contribute to these alterations beyond those that we investigated here.

## Conclusions

We developed an EHT system that can recapitulate key contractile aspects of iron overload cardiomyopathy in human cells. We propose that our in vitro approach could be extended for future basic science and translational studies.

## Acknowledgements

The authors would like to acknowledge the financial support provided by the National Institutes of Health (R01 HL141086 and R01 HL174866 to M.J.G.), the Children’s Discovery Institute of Washington University and St. Louis Children’s Hospital (PM-LI-2019-829 M.J.G.), and the American Heart Association (20CSA35310354 to M.J.G and 970198 to M.J.G. and N.H.).

## Author Contributions

Conception and oversight by N.H. and M.J.G. Cellular and engineered heart tissue studies conducted by S.S.M. with early help from W.T.S. In vitro motility experiments were conducted by L.G. and A.E.G. All authors contributed to the analysis of the data. The first draft was written by S.S.M. and M.J.G. All authors contributed to the writing and/or editing of the manuscript.

## Materials and Methods

### Stem cell culture

Stem cells were cultured as we have described previously [35]. Induced pluripotent stem cells (iPSC) were previously generated from the human BJ fibroblast line (CRL-2522, ATCC) by the Genome Engineering and iPSC Center at Washington University. Stem cells were negative for mycoplasma and had normal karyotypes. Cells were cultured in StemFlex medium (Gibco, Cat # A3349401) in 6 well plates pre-coated with Geltrex (Gibco, Cat # A1413201). Cells were split every 3 or 4 days using 0.02% EDTA (Sigma Aldrich, Cat # E8008).

### Directed differentiation of stem cells to cardiomyocytes and cardiac fibroblasts

Stem cells were differentiated to cardiac cell types using small molecule modulation of Wnt signaling in monolayer culture as we have previously done (**Fig. S1**) [24,25,27,87,88]. Briefly, on days −2 to 0, stem cells were grown in monolayer culture on pre-coated 12-well tissue culture treated plates at density of 5-6 x 10^5^ cells/well in the presence of 1 mL StemFlex with 5 µM ROCK inhibitor, Y27632 (Selleckchem, Cat # S1049). On day 0, the media was changed to contain 7 µM CHIR99021 (Selleckchem, Cat # S1263) in RPMI-1640 medium (Gibco, Cat # 11875135) containing B27 minus insulin supplement (Gibco, Cat # A1895602). On day 3, media was changed to 5 µM IWP2 (Selleckchem, Cat # S7085) in RPMI-1640 medium containing B27 minus insulin supplement. At this point, the protocol changes to derive CMs or CFs.

To generate CMs, cells were cultured in RPMI-1640 medium containing 1x B27 supplement (Gibco, Cat # 17504001) [24,25]. Spontaneous beating of cardiomyocytes was typically observed around day 7. CMs were enriched by culturing in no glucose medium (Gibco, Cat # 11879020) containing 1x B27 Supplement and 4 mM lactate (Sigma, Cat #L4263) on days 12-18. Using this protocol, we regularly achieve > 90% pure cardiomyocytes. All experiments were conducted after day 30.

To generate CFs [27], cells were split on day 5 and cultured for three days in Advanced DMEM/F12 medium (Life Technologies, Cat # 12634-028) containing 5 μM CHIR99021 and 2 μM retinoic acid (Sigma, Cat # R2625). On day 12, cells were changed into Fibroblast Growth Medium 3 (PromoCell, Cat # C23025) containing 10 μM recombinant human FGF2 (R&D Systems, cat # 233-FB) and 10 μM SB431542 (Tocris Bioscience, cat # 1614). Using this procedure, we achieve >90% pure CF [89].

### Iron treatment of cells/tissues

Unless otherwise noted, cells were treated with iron by adding 100 μM ammonium iron (III) citrate (Sigma-Aldrich, Cat # F5879-100G) and 1 mM L-ascorbic acid (Sigma-Aldrich, Cat # A5960-25G) for 5 days prior to experimental measurements.

### Cell viability assays

50,000 cells/well were seeded into 4-well dishes. Cells were cultured in RPMI with insulin medium with or without iron for 5 days. Media for iron treated cells contained ammonium iron (III) citrate (Sigma-Aldrich, Cat # F5879-100G) along with 1 mM L-ascorbic acid (Sigma-Aldrich, Cat # A5960-25G). After 5 days, cells were stained with 1 µM calcein-AM dye (Thermo Fischer, Cat # C3100MP), 1 µg/mL hoechst 33342 (Thermo Fischer, Cat # 62249), and 2 mM EthD-1 (Thermo Fischer, Cat # E1169) in HBSS buffer for 30 minutes at 37 °C. Cells were then washed 3x with HBSS buffer. Fluorescence was measured on an Olympus IX-70 epifluorescence microscope. Cell viability counts were performed in ImageJ [90] across multiple fields of view.

### Measurement of relative intracellular iron levels

Intracellular iron levels were measured using a colorimetric iron kit (abcam, Cat# ab83366) according to the kit directions. Briefly, 5×10^6^ cells were pelleted and lysed in 100 µL of iron assay buffer. Cells were centrifuged at 16,000 x g for 10 minutes at 4°C and the pellet was discarded. 50 µL of supernatant was then used according to the kit instructions. Absorbance was measured using a BioTEK Synergy H1 microplate reader.

### Mechanical testing of ring-shaped tissues

Ring shaped tissues were generated as previously described [91]. Briefly, CMs (1.1 x 10^6^ cells) and CFs (30,000 cells) were mixed with the following solution made on ice: rat collagen I (Corning: 354236; 1 mg/mL final concentration) was combined with equal parts 2x DMEM and the pH was neutralized using 0.5 M Sodium Hydroxide (Sigma, Cat# S5881), 0.77 mg/ml Matrigel, 5 µM ROCK inhibitor, Y27632, seeded into circular PDMS molds, and supplemented with in RPMI-1640 media with 20% FBS, 100 U/mL penicillin-streptomycin, and 10 µM ROCK inhibitor, Y27632 as previously described [19]. Tissues were maintained in culture for 14 days in EHT medium comprised of DMEM (Gibco, Cat # 11965092), 10% FBS (Gibco, Cat # A5256701), 1% MEM non-essential amino acids (Gibco, Cat # 11140050), 1% GlutaMAX (Gibco, Cat # 35050079), and 1% Penicillin-Streptomycin [41].

For mechanical testing, ring-shaped tissues were mounted between a force transducer and a length mover of an Aurora Scientific Instrument (801C: Small Intact Muscle Apparatus). Tissues were preconditioned by being held at a constant force of 2.5 mN for 3 min under 1 Hz stimulation. The tissue’s original length, L_0_, was defined as the tissue length right before force was exerted on the force transducer. The mechanical testing consisted of a series of 8 stretches, increasing the length by 2.5% of L_0_ with each stretch (**Fig. S3**). Data were recorded at 100 Hz and analyzed using custom software in MATLAB. The active and passive forces were calculated as the average of the last five beats before the length change (see **Fig. 2a**). Contraction/relaxation velocities were defined as the peak values calculated from the derivative (**Fig. S4**). Time to 50, 75, and 90% of contractions were calculated as the time from the stimulus to a defined threshold. The time to 50, 75, and 90% of relaxation were calculated as the time from peak force to the defined threshold (**Fig. S4**). Force integrals were defined as the area under the force transient (**Figs. S6a and S6b**).

### Micro–heart muscle (μHM) for fluorescence-based assays

Micro–heart muscles were constructed as previously described [28]. Importantly, these tissues are thin and formed over a cell culture surface to facilitate fluorescence studies. Dog-bone shaped PDMS stencils generated by double-molding a 3D printed template using agar as an intermediary. Stencils were passivated against cell adhesion by treating with a 1:2 dilution of 10% Pluronic F-68 Buffer (Gibco, Cat # 24040-032) in PBS. Tissue culture dishes (GenClone, Cat # 25-200) were coated with 20 µg/mL fibronectin (Sigma-Aldrich, Cat # F1141-5MG) to facilitate cell-attachment. The stencils were gently applied to the tissue culture dish surface in the presence of methanol, before baking in a dry oven at 60 °C. After disinfection with 70% ethanol, the dishes were seeded with CMs (210,000 cells) in RPMI-1640 media with 20% FBS, 100 U/mL penicillin-streptomycin, and 10 µM ROCK inhibitor, Y27632. 24 hours after seeding, media was replaced with RPMI-1640 medium containing 1x B27 supplement with insulin and 100 U/mL Penicillin-Streptomycin. µHMs were then grown for 14 days under uniaxial tension.

### Measurements of calcium transients

Calcium transients were measured using the calcium-sensitive fluorescence reporter FluoForte according to the manufacturer’s instructions (Enzo Life Sciences, Cat # ENZ-52014). μHMs were loaded with 15 µM FluoForte in Tyrode’s solution for 20 minutes at 37°C. Tissues were then washed 3x with Tyrode’s solution before imaging. Fluorescence transients were recorded at 100 fps using a Hamamatsu Orca sCMOS Camera on an Olympus IX-70 microscope while tissues were electrically stimulated at 1 Hz. Calcium transients were analyzed using open-source MATLAB code [18,21].

### Action Potential Waveform Studies

Action potential waveforms measured using the voltage-sensitive fluorescence reporter FluoVolt Membrane Potential kit according to manufacturer’s instructions (Thermo Fischer, Cat # F10488). μHMs were loaded with 2 ml FluoVolt loading solution (1:10 dilution of component A into component B) in Tyrode’s solution for 20 minutes at 37°C. Tissues were then washed 3x with Tyrode’s solution and 200 µl Neuro Background Suppressor (Component C) was added to the buffer before imaging. Fluorescence transients were recorded at 100 fps using a Hamamatsu Orca sCMOS Camera on an Olympus IX-70 microscope while tissues were electrically stimulated at 1 Hz. Action potential waveform morphology was analyzed using open-source MATLAB code [18].

### Reactive oxygen species (ROS) measurements

ROS levels were measured using the ROS-Glo H_2_O_2_ kit according to manufacturer’s instructions (Promega, Cat # G8820). 80,000 cells/well in RPMI with insulin medium were seeded into 96 well plates. On the day of the assay, 25 µM H_2_O_2_ substrate solution at total volume of 100 µL was added to the cells for 6 hours at 37 °C. An equal volume of ROS-Glo Detection Solution was then added and incubated for 20 minutes at room temperature before bioluminescence was read on a BioTEK Synergy H1 plate reader.

### In vitro motility assays

In vitro actin filament gliding assays were performed as previously described [35]. Porcine cardiac actin and myosin were pretreated with iron on ice prior to the experiments with or without iron in the final assay buffer. Myosin was attached to nitrocellulose coated surfaces, the surface was blocked with BSA, and actin filaments were added in the presence of ATP as previously described [35]. Videos were collected at 1-10 seconds per frame depending on the motile rate. Motile rate was quantified by manual tracking using the MTrackJ in ImageJ as we have previously done [35].

### Statistical Analysis

All cellular experiments included at least three technical and biological (differentiations) replicates. All experiments were conducted in parallel with controls on the same day. Statistical testing was done as described in the text using GraphPad Prism. Shapiro-Wilk tests were used to test for normalcy. ANOVAs were conducted where appropriate followed by post-hoc tests with appropriate correction for multiple comparisons. For normally distributed data, 2-tailed t-tests were used, and multiple comparisons were corrected using Tukey’s HSD test. For non-parametric data, we used Mann-Whitney tests followed by Dunn’s correction.

