## Supplement for "Molecular and Cellular Determinants of Human Iron Overload Cardiomyopathy"

### Supplementary Figures

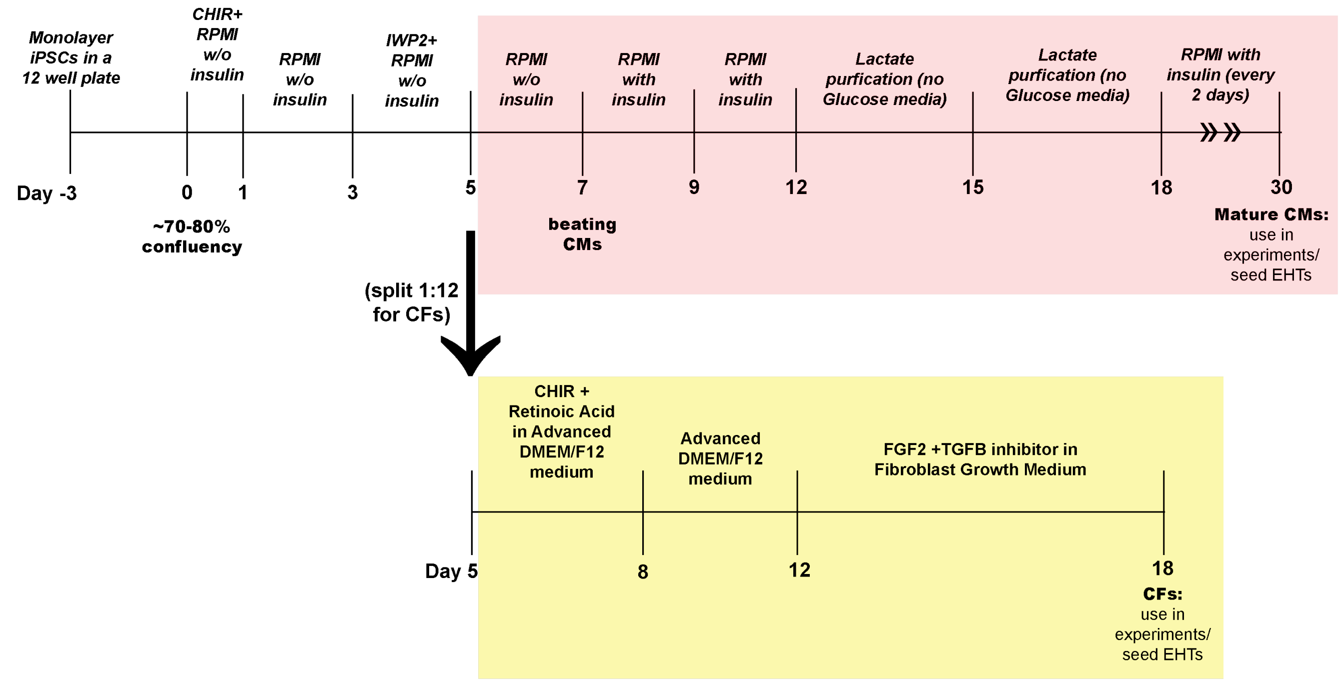


**Figure S1: Experimental timeline for CM and CF differentiation**

Both CMs and CFs were derived from iPSC as previously described . Similar procedures were followed until day 5. The CM protocol is shown in pink, and the CF protocol is shown in yellow.

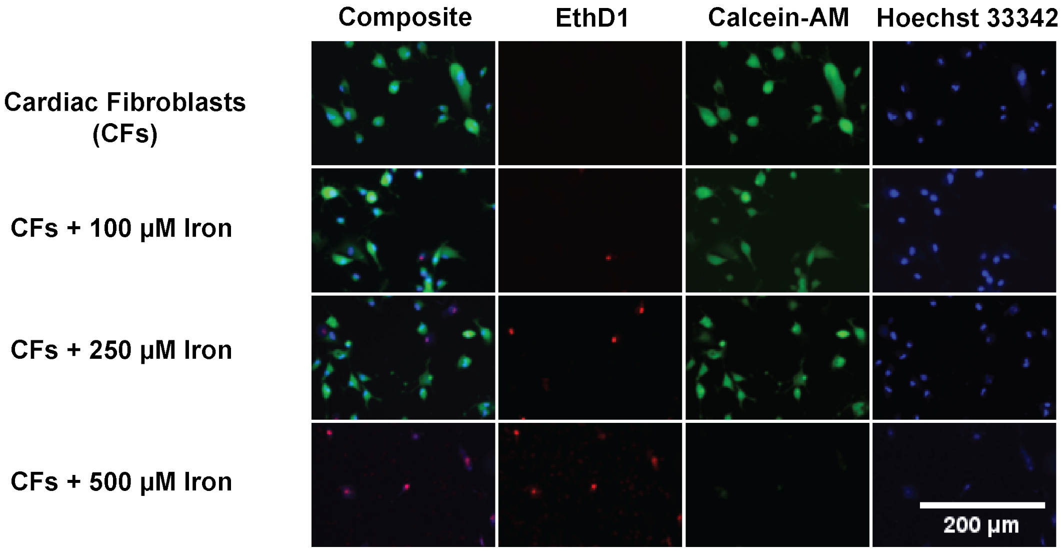


**<u>Figure S2: Viability of cardiac fibroblasts (CFs) in monolayer culture in response to iron.</u>**

Representative images of CFs survival after 5 days of iron treatment analyzed by cell staining with Calcein-AM (green), Hoechst 33343 (blue) and EthD1 (red). Quantification can be found in **Fig. 1c**.

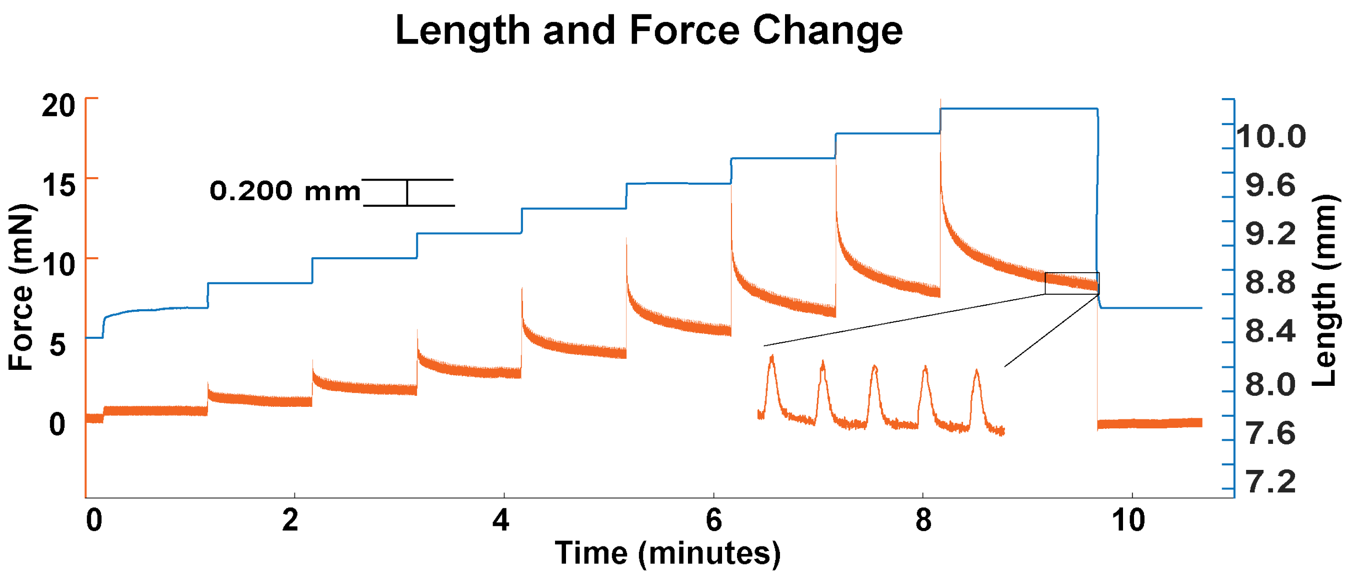


**Figure S3: Representative data trace showing mechanical testing of a ring-shaped tissue.**

A ring-shaped tissue was mounted between a length mover and a force transducer and stimulated at 1 Hz. The data trace shows the force (orange) and the length of the tissue (blue) as a function of time. The tissue was stretched by increments of 2.5% of the initial length at each length step. Inset shows beating of the tissue.

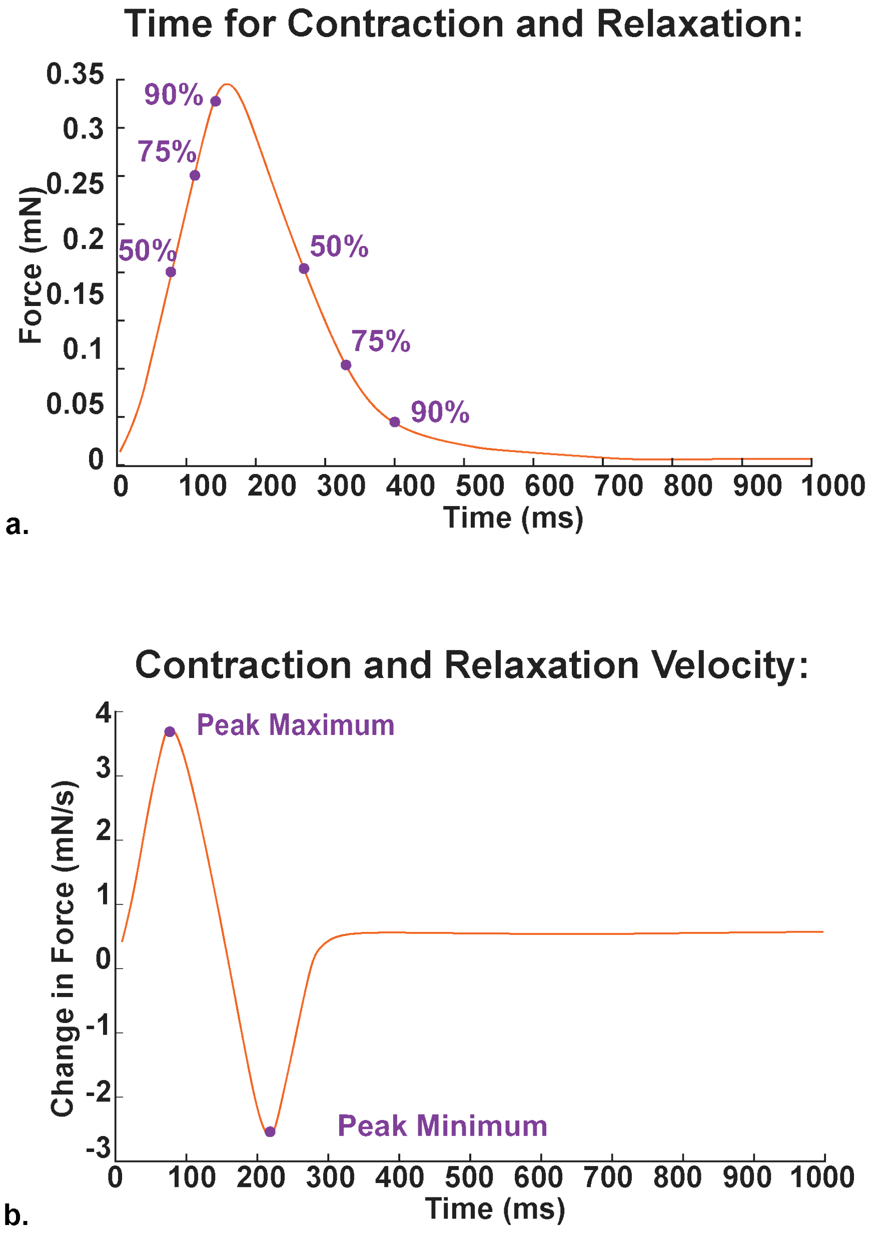


**Figure S4: Calculation of the kinetics of contraction and relaxation.**

Idealized traces showing (a) the active force generated by a tissue as a function of time and (b) the change in force as a function of time. Key metrics reported in the paper are labeled on these traces.

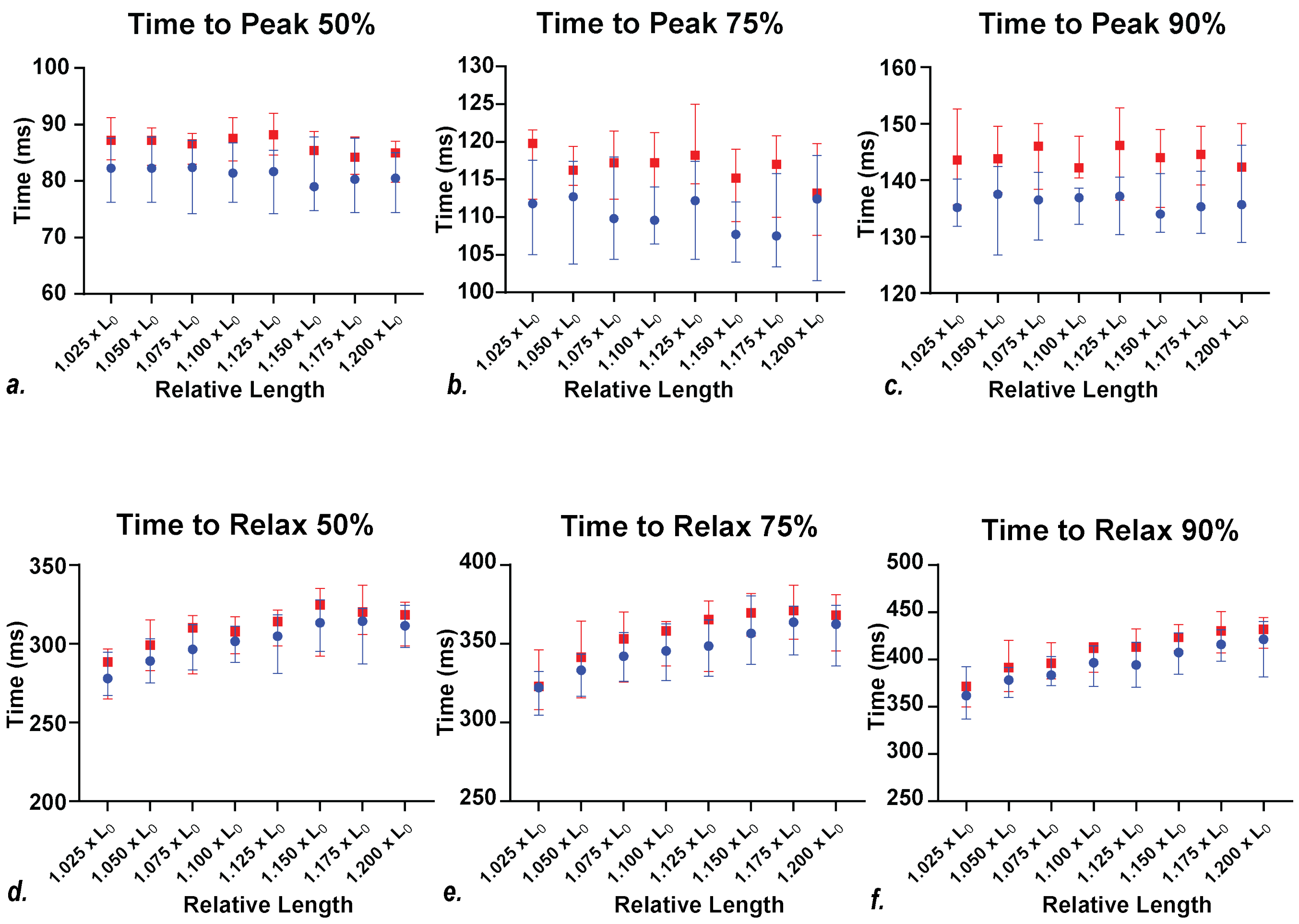


**Figure S5: Kinetics of ring tissue contraction measured by the force transducer.**

Time to peak (TTP) and time to relax (TTR) both were found to be different between control and iron tissues. (TTP: 50% - p-value = 0.0013, 75% and 90% - p-value < 0.001, TTR: 50% - p-value = 0.01, 75% - p-value = 0.02, 90% - p-value = 0.01). Time to relax was also different as function of stretch (TTR: 50% - p-value = 0.0004, 75% and 90% - p-value < 0.0001). n = 24 control tissues (blue) and n = 25 iron-treated tissues (red). Symbols show median and error bars show 95% confidence intervals. Statistical testing was done by two-way ANOVA.

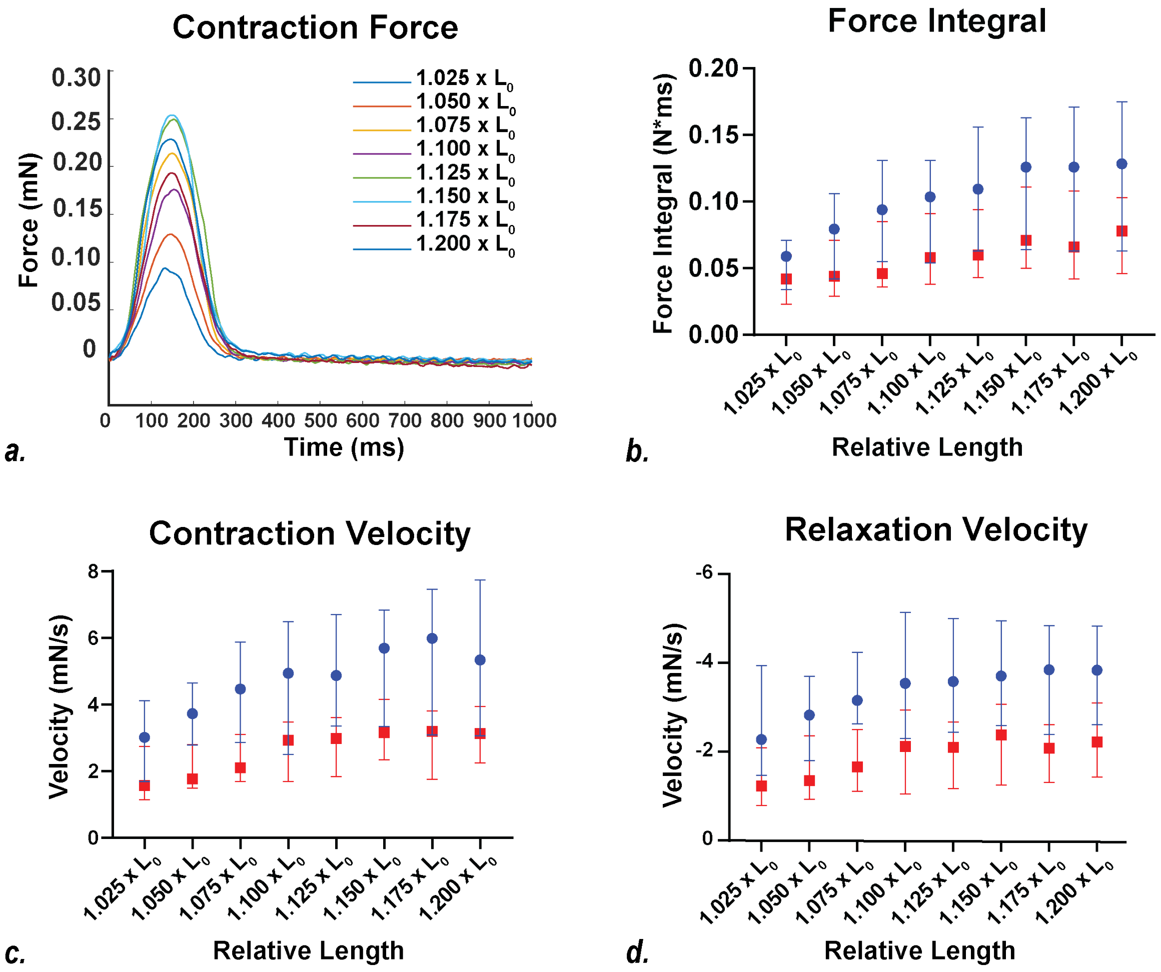


**Figure S6: Key measured parameters for ring tissues mounted on a force transducer.**

(a.) Representative trace showing a single averaged beat for the same tissue are shown as a function of stretch. (b-d) n = 24 control tissues (blue) and n = 25 iron-treated tissues (red). Symbols show median and error bars show 95% confidence intervals. Statistical testing was done by two-way ANOVA. (b.) Force integral (as function of stretch: p-value $<0.0001$ and control vs iron: p-value $< 0.0001$), (c.) contraction velocity (as function of stretch: p-value $<0.0001$ and control vs iron: p-value $< 0.0001$), and (d.) relaxation velocity (as function of stretch: p-value = 0.004 and control vs iron: p-value $< 0.0001$) are shown.

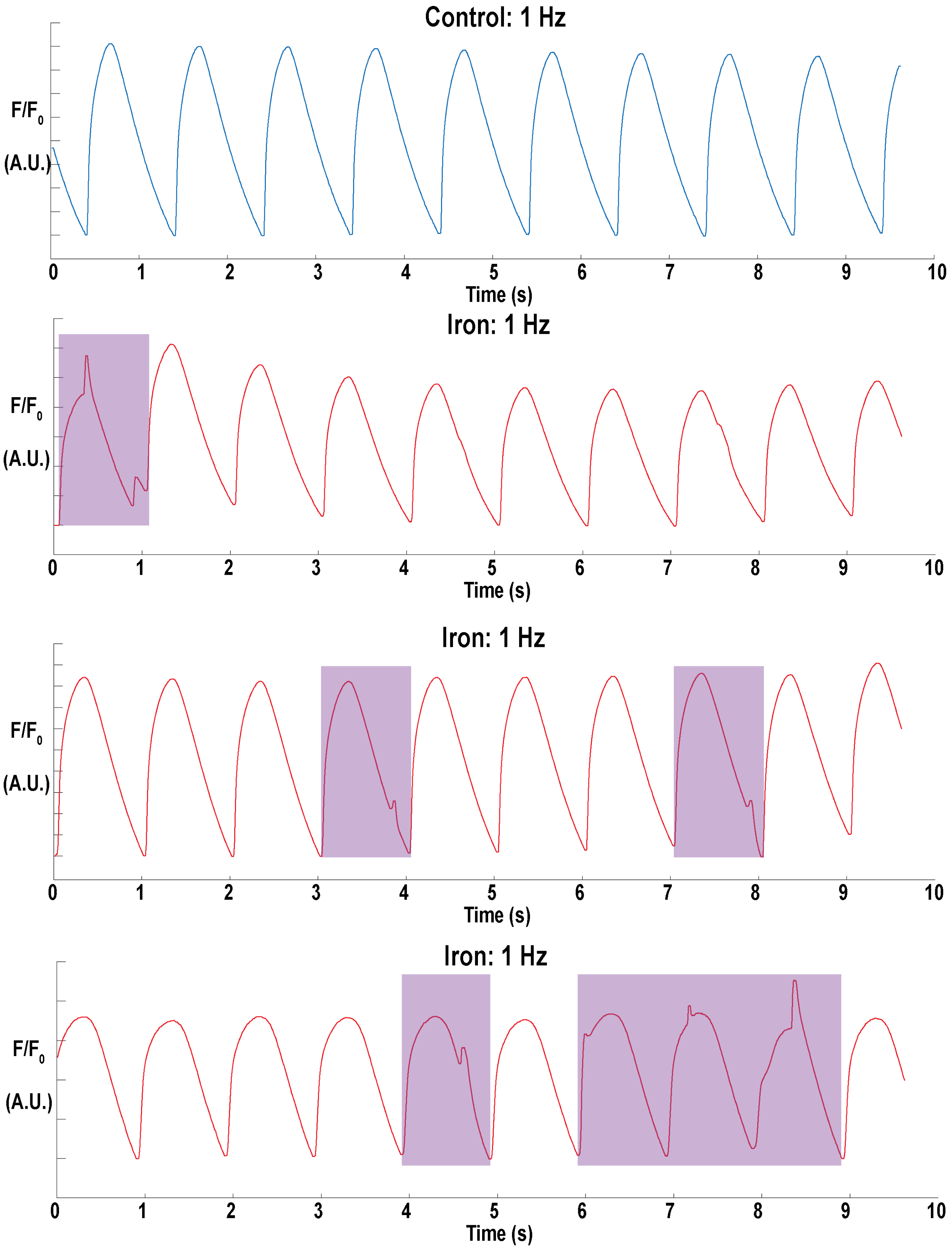

Control: 1 Hz
F/F0
(A.U.)
0 1 2 3 4 5 6 7 8 9 10
Time (s)
Iron: 1 Hz
F/F0
(A.U.)
0 1 2 3 4 5 6 7 8 9 10
Time (s)
Iron: 1 Hz
F/F0
(A.U.)
0 1 2 3 4 5 6 7 8 9 10
Time (s)
Iron: 1 Hz
F/F0
(A.U.)
0 1 2 3 4 5 6 7 8 9 10
Time (s)

**Figure S7: Representative calcium waveforms for µHMs loaded with a fluorescent calcium indicator.**

Waveforms were collected under 1 Hz stimulation. Aberrant waveforms are colored with purple boxes. Quantification can be found in **Fig. S10**.

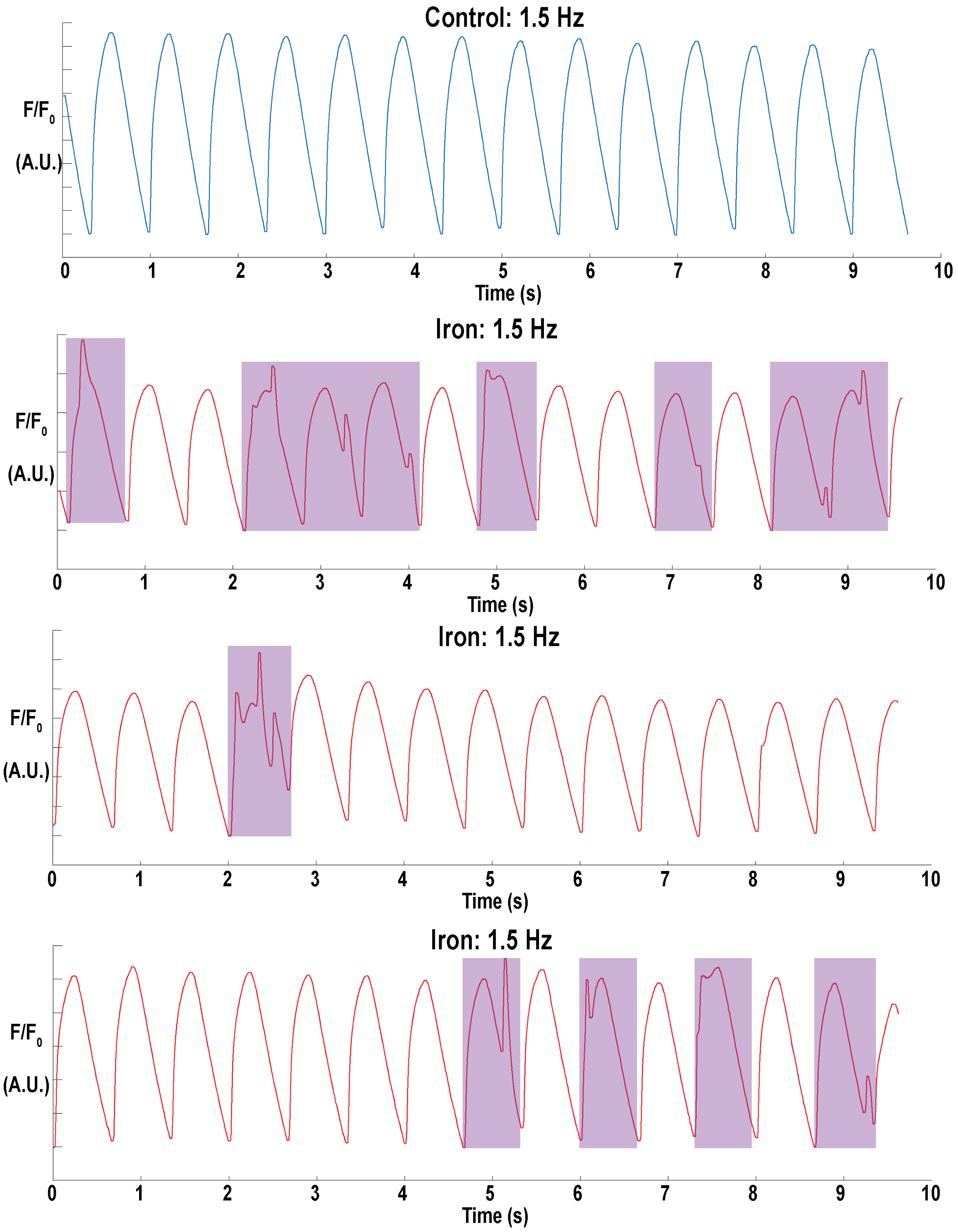

Control: 1.5 Hz
F/F0
(A.U.)
0
1
2
3
4
5
6
7
8
9
10
Time (s)
Iron: 1.5 Hz
F/F0
(A.U.)
0
1
2
3
4
5
6
7
8
9
10
Time (s)
Iron: 1.5 Hz
F/F0
(A.U.)
0
1
2
3
4
5
6
7
8
9
10
Time (s)
Iron: 1.5 Hz
F/F0
(A.U.)
0
1
2
3
4
5
6
7
8
9
10
Time (s)

**Figure S8: Representative calcium waveforms for µHMs loaded with a fluorescent calcium indicator.**

Waveforms were collected under 1.5 Hz stimulation. Aberrant waveforms are colored with purple boxes. Quantification can be found in **Fig. S10**.

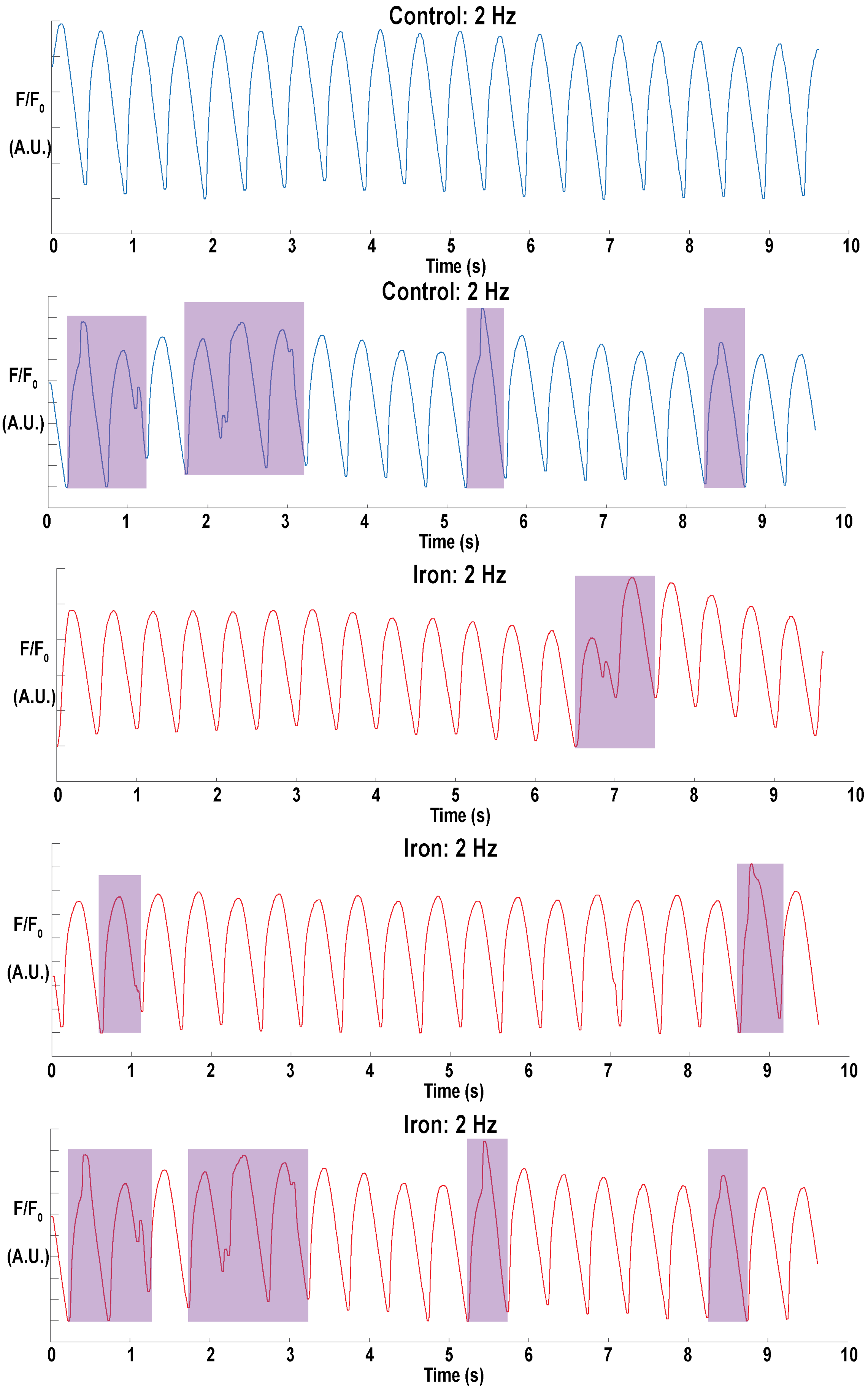
Control: 2 Hz
F/F0
(A.U.)
0 1 2 3 4 5 6 7 8 9 10
Time (s)
Control: 2 Hz
F/F0
(A.U.)
0 1 2 3 4 5 6 7 8 9 10
Time (s)
Iron: 2 Hz
F/F0
(A.U.)
0 1 2 3 4 5 6 7 8 9 10
Time (s)
Iron: 2 Hz
F/F0
(A.U.)
0 1 2 3 4 5 6 7 8 9 10
Time (s)
Iron: 2 Hz
F/F0
(A.U.)
0 1 2 3 4 5 6 7 8 9 10
Time (s)

**Figure S9: Representative calcium waveforms for µHMs loaded with a fluorescent calcium indicator.**

Waveforms were collected under 1 Hz stimulation. Aberrant waveforms are colored with purple boxes. Quantification can be found in **Fig. S10**.

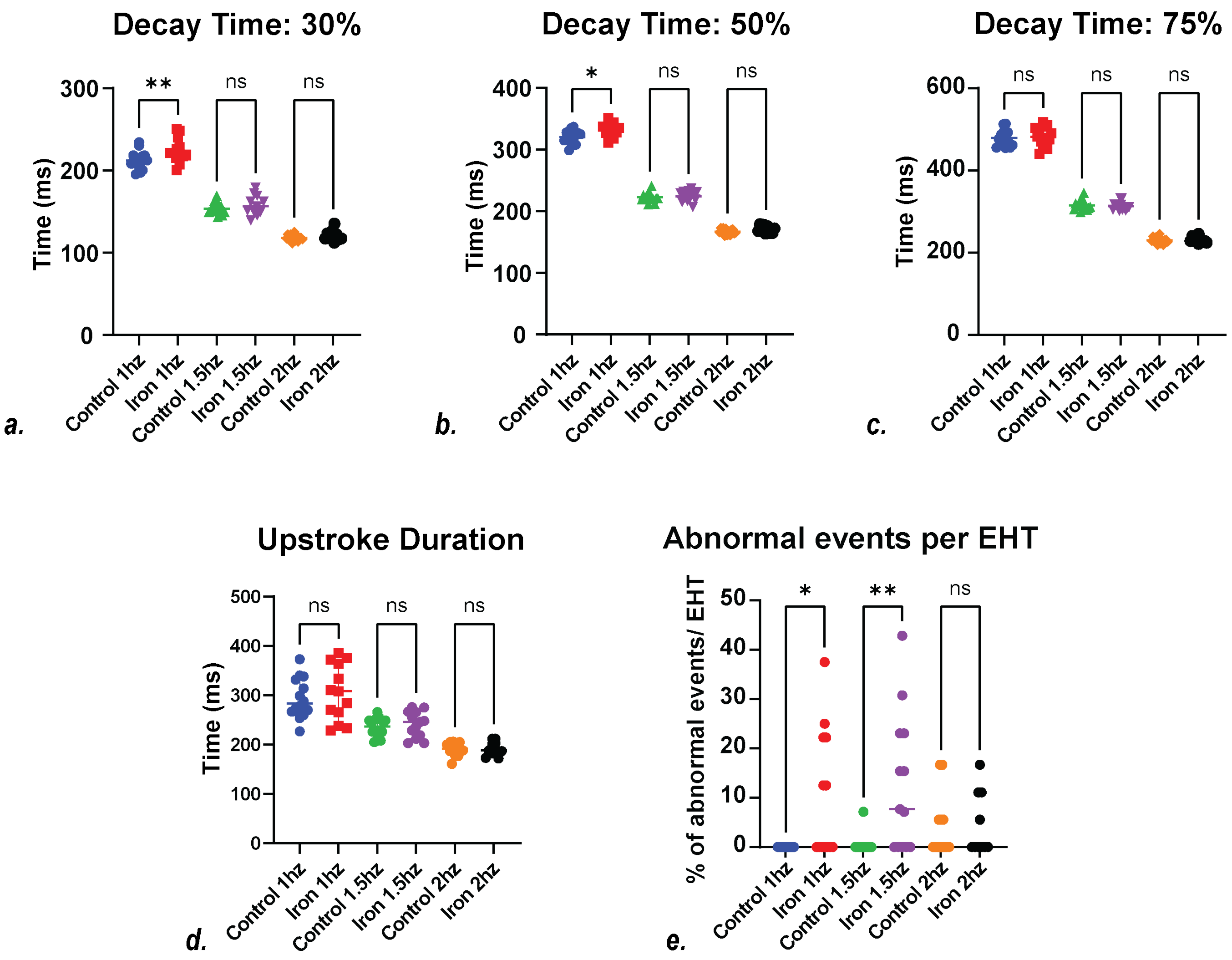


**Figure S10: Effect of iron on calcium transients in μHMs paced at different frequencies.**

Key parameters were measured for tissues paced at 1 Hz; n = 14 control and n = 13 iron fed tissues, 1.5 Hz; n = 14 control and n = 13 iron fed tissues, and 2 Hz; n = 13 control and n = 10 iron fed tissues. Shown are the time for the force to decay by (a.) 30, (b.) 50, and (c.) 75%. (d.) Upstroke duration. Only decay values for control and iron treated tissues stimulated at 1 Hz were statistically different; p-value < 0.001 Statistical testing was done by ordinary one-way ANOVA, and multiple comparisons were corrected by Tukey's tests (e.) The percentage of abnormal calcium waveforms seen per 10 seconds. Statistical testing was done by Kruskal-Wallis test. *p-value≤0.05, **p-value≤0.01, ***p-value≤0.01, and ****p-value≤0.01.

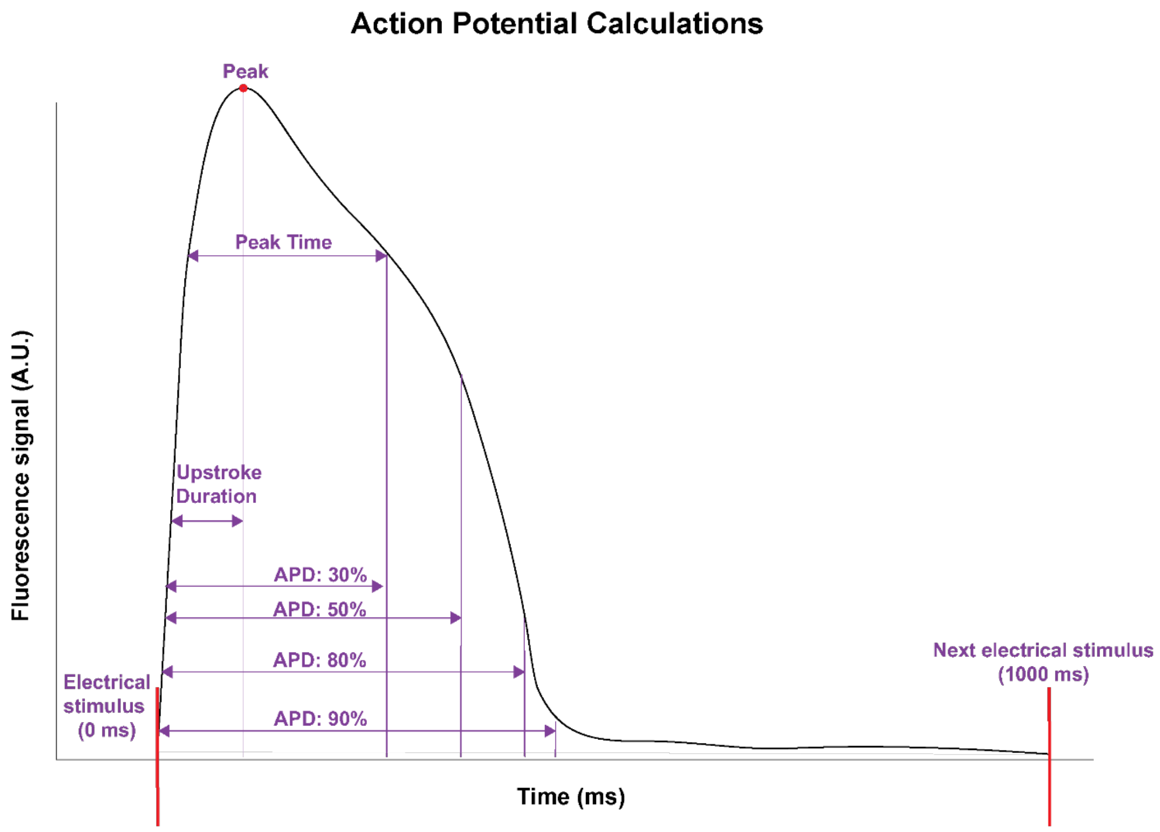


**Figure S11: Calculation of the kinetics of the action potential.**

Idealized traces showing the action potential generated by a tissue as a function of time. Key metrics reported in the paper are labeled on these traces.

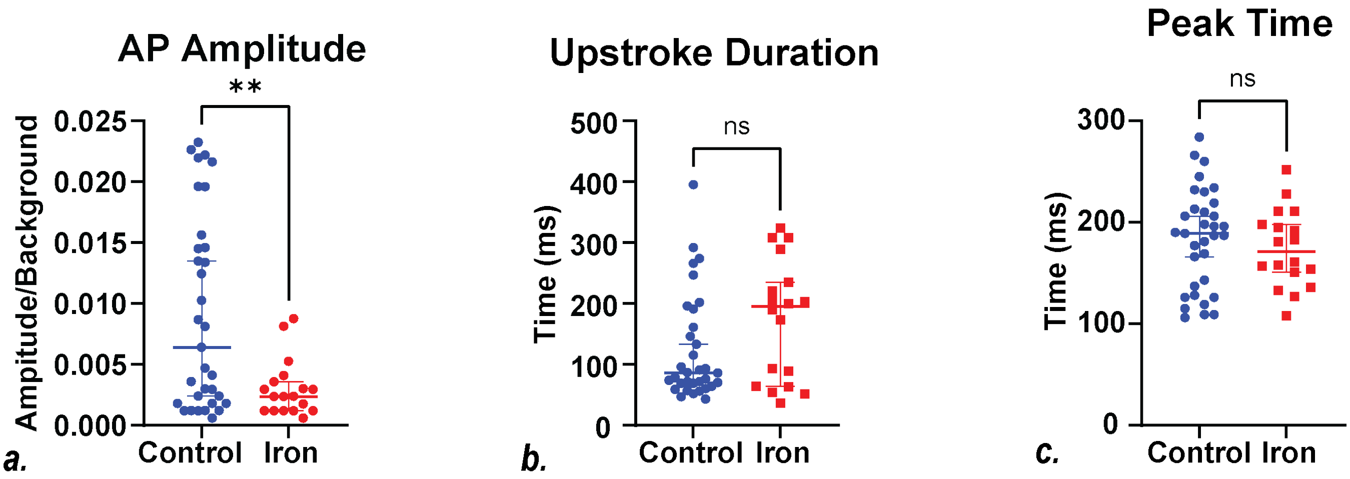


**<u>Figure S12: Key parameters measured for the action potentials of µHMs paced at 1 Hz.</u>**

Action potentials waveforms were examined from n = 33 control (blue) and n = 18 iron fed (red) µHMs. Shown are the (a.) action potential (AP) amplitude, (b.) upstroke duration, and (c.) peak time. Symbol shows median and the error bars show 95% confidence intervals. Statistical testing was done using a Mann-Whitney test. **p-value≤0.01.

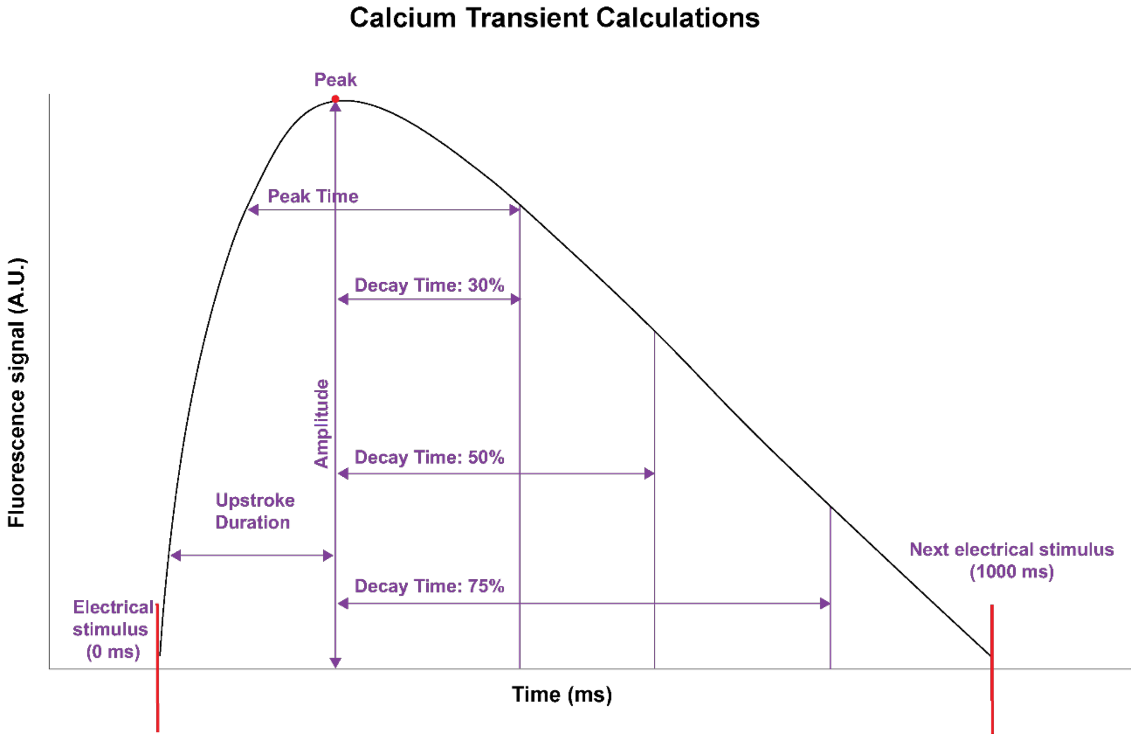


**<u>Figure S13: Calculation of the kinetics of the calcium transient.</u>**

Idealized traces showing the calcium transient generated by a tissue as a function of time. Key metrics reported in the paper are labeled on these traces.

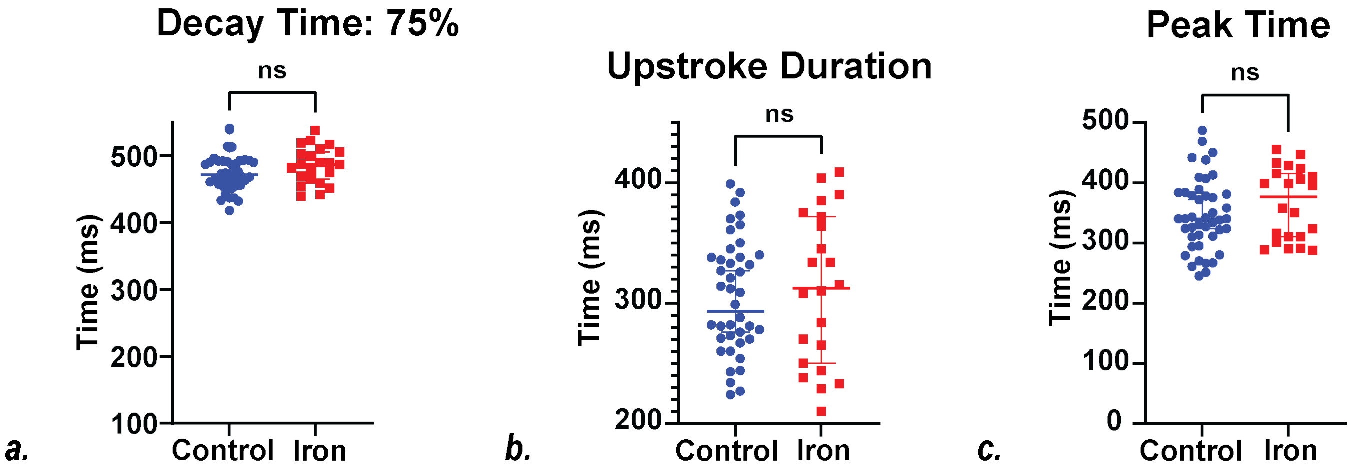


**<u>Figure S14: Effect of iron on the kinetics of the calcium transient in µHMs paced at 1 Hz.</u>**

Calcium transient waveforms were examined from n = 44 control (blue) and n = 22 iron fed (red) µHMs. Shown are the (a.) time to decay by 75%, (b.) upstroke duration, and (c.) peak time. Central bar shows the median and the error bars show 95% confidence intervals. Statistical testing was done using a Mann-Whitney test.

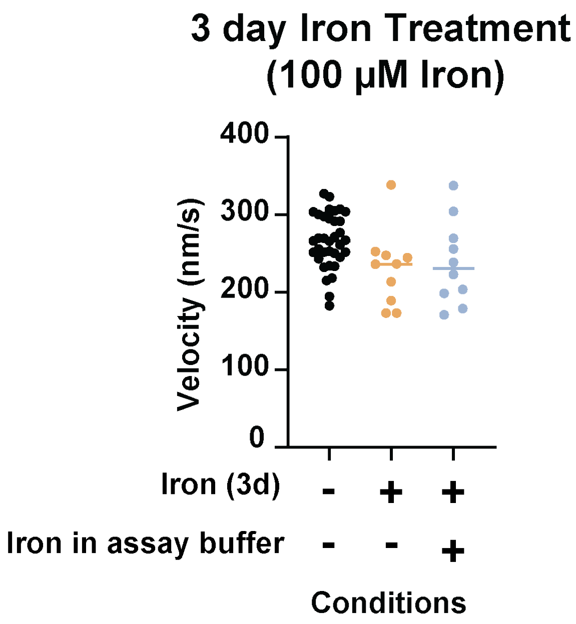


**Figure S15: Additional *in vitro* motility assays investigating iron effects on the actomyosin contractile apparatus.**

Motile rate of actin filaments moving over a bed of cardiac myosin. Each point represents an individual actin filament speed recorded over 30 seconds. Statistical testing was done using an ANOVA followed by post-hoc t-tests.
